# aCPSF1 controlled archaeal transcription termination: a prototypical eukaryotic model

**DOI:** 10.1101/843821

**Authors:** Lei Yue, Jie Li, Bing Zhang, Lei Qi, Fangqing Zhao, Lingyan Li, Xiuzhu Dong

## Abstract

Transcription termination defines RNA 3′-ends and guarantees programmed transcriptomes, thus is an essential biological process for life. However, transcription termination mechanisms remain almost unknown in Archaea. Here reported the first general transcription termination factor of Archaea, the conserved ribonuclease aCPSF1, and elucidated its 3′-end cleavage dependent termination mechanism. Depletion of *Mmp-aCPSF1* in a methanoarchaeon *Methanococcus maripaludis* caused a genome-wide transcription termination defect and overall transcriptome chaos, and cold-sensitive growth. Transcript-3′end-sequencing (Term-seq) revealed transcriptions mostly terminated downstream of a uridine-rich terminator motif, where *Mmp*-aCPSF1 performed cleavage. The endoribonuclease activity was determined essential to terminate transcription *in vivo* as well. Through super-resolution photoactivated localization microscopy imaging, co-immunoprecipitation, and chromatin immunoprecipitation, we demonstrated that *Mmp*-aCPSF1 localizes within nucleoid and associates with RNAP and chromosomes. aCPSF1 appears to co-evolve with archaeal RNAPs, and two distant orthologs each from *Lokiarchaeota* and *Thaumarchaeota* could replace *Mmp*-aCPSF1 to termination transcription. Thus, aCPSF1 dependent termination mechanism could be universally employed in Archaea, including *Lokiarchaeota*, one supposed archaeal ancestor of Eukaryotes. Therefore, the reported aCPSF1 cleavage-dependent termination mode not only hints an archetype of Eukaryotic 3′-end processing/cleavage triggered RNAP II termination, but also would shed lights on understanding the complex eukaryotic termination based on the simplified archaeal model.

## Introduction

Transcription, the fundamental biological process in transforming genetic information from DNA to proteins, must be properly terminated^1–3^. Transcription termination functions in i) defining RNA 3′-ends, ii) preventing read-through resulted inappropriate expression of downstream genes or antisense transcriptions, iii) recycling RNA polymerase (RNAP), and iv) minimizing biomacromolecular machinery collision^2–4^. Consequently, organisms have evolved precise mechanisms to govern transcription termination^2–5^. Bacteria primarily employ Rho-dependent and -independent (intrinsic) termination mechanisms. The former relies on an RNA translocase, Rho, to dissociate the transcription elongation complex (TEC), while the latter depends merely on nascent RNA structures harboring 7-8 base-paired hairpins followed by adjacent uridine residues^2, 3, 6, 7^. In Eukaryotes, RNAP II, which transcribes mRNAs and non-coding RNAs, employs multifaceted termination strategies by an involvement of transcript 3′-end processing event; the poly(A) sites, one 3′-end processing/termination signal, are recognized by a cleavage and polyadenylation complex (CPF/CPSF), in which the endoribonuclease, human CPSF73 or yeast Ysh1, cleaves RNA downstream and followed by polyadenylation to ensure mRNA maturation and trigger RNAP II dissociation for transcription termination^3, 5^. In addition, other factors are required for efficient dismantling of RNAP II ^3, 5, 8–12^.

However, little is known about transcription termination in Archaea^1, 13^, the third domain of life. Archaea are thought to represent ancient earth life and possess eukaryotic-like genetic information processing systems to express bacteria-like genomic information^1, 14^. Particularly, they employ the eukaryotic RNAP II orthologs, archaeal RNAP (aRNAP)^15^, but have compact genomes with short intergenic-regions (IGRs) and polycistronic operons usually co-transcribed. Therefore, transcription termination in Archaea must be elaborately controlled. Individual gene and genome-level 3′-ends sequencing (Term-seq) analyses have identified representative uridine-rich tracts in archaeal transcripts 3′-ends, but without preceding hairpin structures^1, 13, 16, 17^. Term-seq also identified eukaryotic-like alternative 3′-untranslated regions (3′-UTR) isoforms generated from the consecutive termination sites in two representative archaeal species, *Methanosarcina mazei* and *Sulfolobus acidocaldarius*^13^. This suggests archaea could employ eukaryotic-like transcription termination mechanisms and depend on trans-acting (termination) factors^13^. However, general transcription termination factors have not been discovered in Archaea thus far^1, 13, 18^. Although Eta was reported to release stalled TECs from damaged DNA sites in *Thermococcus kodakarensis*^18^, it functions specially in DNA damage response^1, 14, 18^.

In this work, we revealed that the conserved aCPSF1, depending on its ribonuclease activity, controls the genome-wide transcription termination and ensures programmed transcriptome and the optimal physiology in Archaea, thus is the first reported archaeal general termination factor. aCPSF1 appeared to have co-evolved with archaeal RNAP, and two distant orthologs from *Lokiarchaeota* and *Thaumarchaeota* could substitute *Mmp*-aCPSF1 to implement transcription termination in *Methanococcus*, suggesting that aCPSF1 depended transcription termination could be wildly employed in Archaea. Noticeably, aCPSF1 cleaves downstream the uridine-rich terminator motif to produce transcripts 3′-ends but no polyadenylation, so resembles its eukaryotic orthologs (yeast Ysh1 and human CPSF73) in Eukaryotic RNAP II transcription termination, and could expose an evolutionary relic of eukaryotic transcription termination.

## Results

### Depleted expression of *Mmp*-aCPSF1 causes cold-sensitive growth and disordered transcriptome

Archaeal CPSF1 (aCPSF1) affiliates within the β-CASP family of ribonucleases, and *in vitro* assays have identified three aCPSF1 orthologs exhibiting endoribonuclease activities^19–21^ along with one also displaying 5′-3′ exoribonuclease activity^20^. aCPSF1 is universally distributed in all defined archaeal phyla^20^ and determined as an essential gene by Sarmiento *et al.*^22^ and our repeated failures in deleting the encoding gene in *Methanococcus maripaludis.* These highlight its fundamental role in Archaea. To investigate its function, we constructed an expression depleted strain of *Mmp*-aCPSF1 (▽*aCPSF1*) in *M. maripaludis* using a *TetR*-*tetO* repressor-operator system (Fig. 1a). Compared with wild-type strain S2, only 60% and 20% *Mmp*-aCPSF1 abundances were determined in ▽*aCPSF1* (minus tetracycline) when grown at 37°C and 22°C, respectively, indicating a successful depletion of *Mmp-aCPSF1* in ▽*aCPSF1* (Fig. 1b and Supplementary Fig. 1a). Although ▽*aCPSF1* had a similar growth rate as strain S2 at 37°C (Supplementary Fig. 1b), it exhibited markedly reduced growth at 22°C (*μ*=0.035 vs 0.06 h^-1^ of S2, Fig. 1b). These coincide with the three-fold lower *Mmp*-aCPSF1 protein in ▽*aCPSF1* grown at 22°C than at 37°C. Concomitantly, ∼2-fold extended half-life was observed for 22°C-cultured ▽*aCPSF1* than that of S2 (8.4 min vs 4.7 min, Fig. 1c). These indicated that aCPSF1 is an archaeal house-keeping ribonuclease functioning in mRNA turnover.

**Figure 1.**
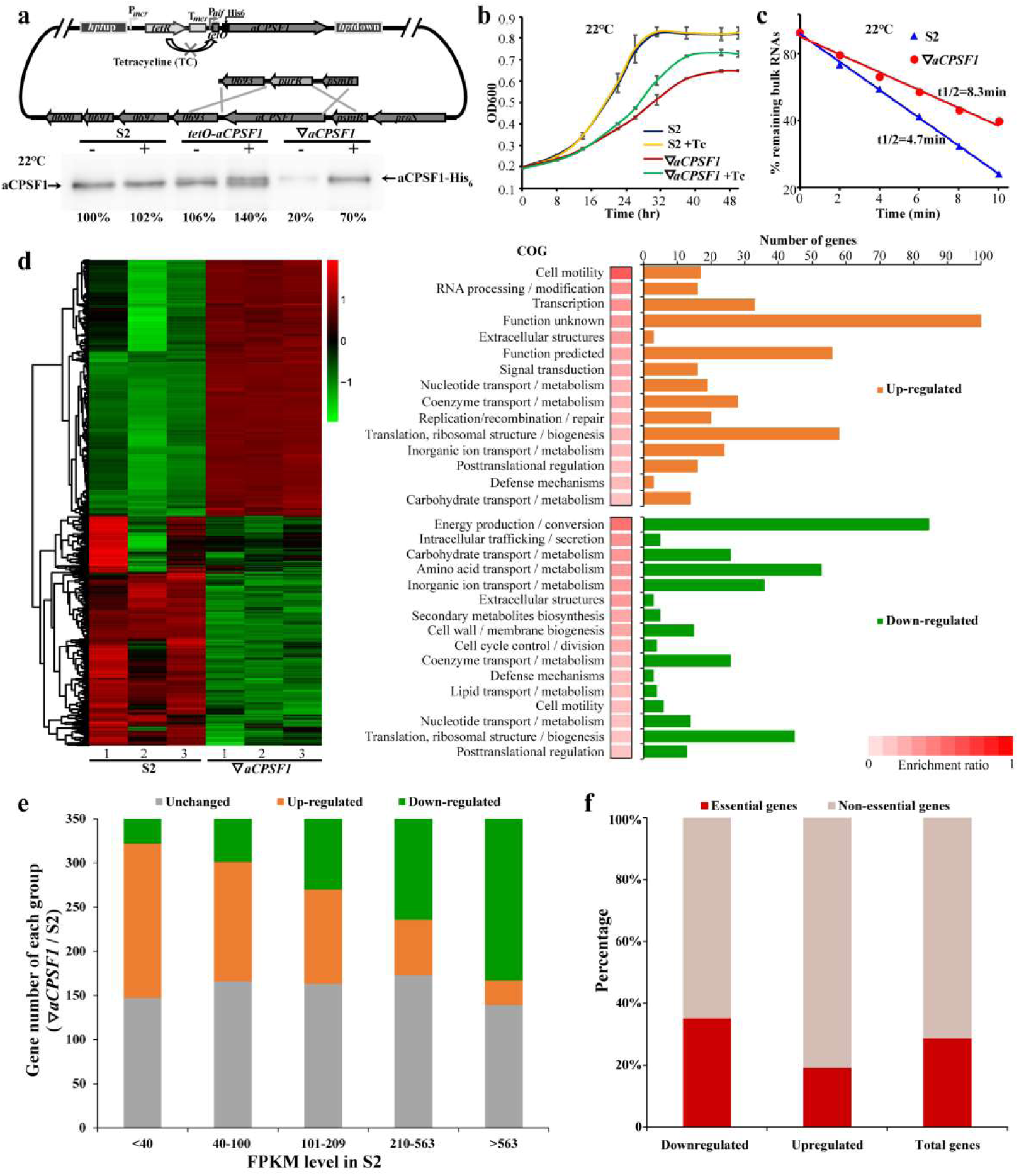
Depleted expression of *Mmp-aCPSF1* caused cold-sensitive growth, prolonged RNA lifespan, and disordered transcriptome in *M. maripaludis*. **a**, A schematic depicting the construction strategy of *Mmp-aCPSF1* (*MMP0694*) depletion strain (▽*aCPSF1*) as described in Methods. The tetracycline (Tc) responsive regulator (*tetR*) and operator (*tetO*) cassette were fused to the His_6_-tagged N-terminus of *Mmp-aCPSF1* and then inserted into the *hpt* locus based on 8-azahypoxanthine resistance selection to obtain an ectopic *Mmp*-*aCPSF1* expression strain (*tetO-aCPSF1*). Next, the indigenous *Mmp-aCPSF1* gene was in-frame replaced with puromycin resistance gene (*pur^R^*) resulting in the TetR regulatory *Mmp*-aCPSF1 depleted strain (▽*aCPSF1*). P*mcr* and T*mcr*, promoter and terminator of methoanococcal methyl-CoM reductase (*mcr*), respectively; P*nif*, the core promoter sequence including BRE and TATA box of the methoanococcal *nif* (nitrogen fixation gene). Lower panel shows western blot detected *Mmp*-aCPSF1 protein abundance (percentages referenced to lane 1) by addition (+) or not (–) of 100 μg/ml tetracycline (Tc) in S2 (wild-type), *tetO-aCPSF1*, and ▽*aCPSF1* at 22°C. The aCPSF1 and aCPSF1-His_6_ labeled flanking the gel pointed the indigenous and *hpt*-site inserted His_6_-*Mmp*-aCPSF1, respectively. **b**,**c**, *Mmp*-*aCPSF1* depletion reduced growth (**b**) and prolonged the bulk mRNA half-lives (**c**) at 22°C. Strains S2 and ▽*aCPSF1* were cultured with or without addition of tetracycline (Tc). Three batches of culture of each strain were measured, and standard deviations were shown. Bulk mRNA half-lives were quantified by the [^3^H]-uridine signal attenuation as described in Methods. **d**, Hierarchical clustering (left) and functional category (right) analysis of the differential transcribed genes in strains S2 and ▽*aCPSF1* grown at 22°C in triplicate. Heat plot representation of Log2 of differential expression ratio was shown with color intensity. Green and red represent the minima and maxima abundance fold, respectively. The functional category enrichment ratio (%) was shown as the gene numbers that account for the up (orange)- or down (green)- regulated genes divided by the total gene numbers in each COG categories, and the top 15 and 16 mostly enriched up- and down-regulated gene categories were shown respectively. **e**, Hierarchical grouping on differential transcribed genes by *Mmp-aCPSF1* depletion. Based on the transcript abundance (FPKM) in strain S2, all the genes were grouped into five ranks (<40, 40-100, 101-209, 210-563, and >563), and then the transcriptional up- and down-regulated, and unchanged gene numbers in ▽*aCPSF1* were counted for each rank. **f**, According to the functional essentiality of genes in *M. maripaludis* defined by Sarmiento et al.^22^, percentages of the essential and non-essential genes in the transcriptional up- and down-regulated, and unchanged gene groups in ▽*aCPSF1* were calculated, respectively.

Differential transcriptomics using strand-specific RNA-seq revealed that *Mmp*-*aCPSF1* depletion altered transcriptions of ∼55% genes (947) at 22°C, but with an unexpected decreased expression of ∼26% genes (453) including many housekeeping genes (Fig. 1d and Supplementary Dataset 1). This contradicts with the ∼2-fold prolonged bulk mRNA life-span in ▽*aCPSF1* and the ribonuclease activity of aCPSF1. By hierarchically grouping all transcripts into five ranks based on their FPKMs in strain S2, we found that the lower expressed transcript ranks in S2 were mostly upregulated in ▽*aCPSF1*, while the higher expressed transcript ranks in S2 were mostly downregulated in ▽*aCPSF1* (Fig. 1e), behaving like a “robbing the rich to feed the poor” pattern caused by *Mmp-aCPSF1* depletion. In addition, expression of ∼40% and ∼20% of the possibly essential genes defined by Sarmiento *et al.*^22^ was reduced and increased in ▽*aCPSF1*, respectively (Fig. 1f). This suggests that *Mmp*-aCPSF1 plays a role in controlling the ordered transcriptome.

### *Mmp-aCPSF1* depletion results in genome-wide transcription read-throughs

Strikingly, querying the strand-specific RNA-seq mapping files found prevalent prolonged transcript 3′-ends in ▽*aCPSF1*. Such as *MMP1100*, a predicated master regulator for methanogenesis^23^, its 3′-end was highly mapped and markedly extended, and resulted antisense transcription of *MMP1099* on opposite strand (Fig. 2a left). 3′-end extension from the upstream *MMP1147* (encoding ribosomal subunit L37) into the downstream *MMP1146* region on same strand was also observed (Fig. 2b left). Northern blot (Fig. 2a, b and Supplementary Fig. 2 middle) and rapid amplification of cDNA 3′-ends (3′RACE) (Fig. 2a, b and Supplementary Fig. 2 right) verified the transcription read-throughs (TRTs) in ▽*aCPSF1*. The *M. maripaludis* genome comprises 1,722 coding genes that are organized into 1,079 predicted transcription units (TUs) with a median IGR of 200 bp (Supplementary Dataset 2). Pair-wisely comparing the FPKMs of a TU and its IGR in S2 and ▽*aCPSF1* at genome-level, and further metaplot visualization by normalization of 750 TUs indicated a genome-wide FPKM increase in IGRs but reduction in TU bodies in ▽*aCPSF1* (Fig. 2c, d and Supplementary Dataset 2). Similarly, statistical comparison for the 1,079 TUs in ▽*aCPSF1* and S2 yielded much higher fold of transcriptional increases in IGRs (median: 4.36) than in TU bodies (median: 1.21) (Fig. 2e and Supplementary Fig. 3). Thus, *Mmp*-*aCPSF1* depletion caused genome-wide TRTs.

**Figure 2.**
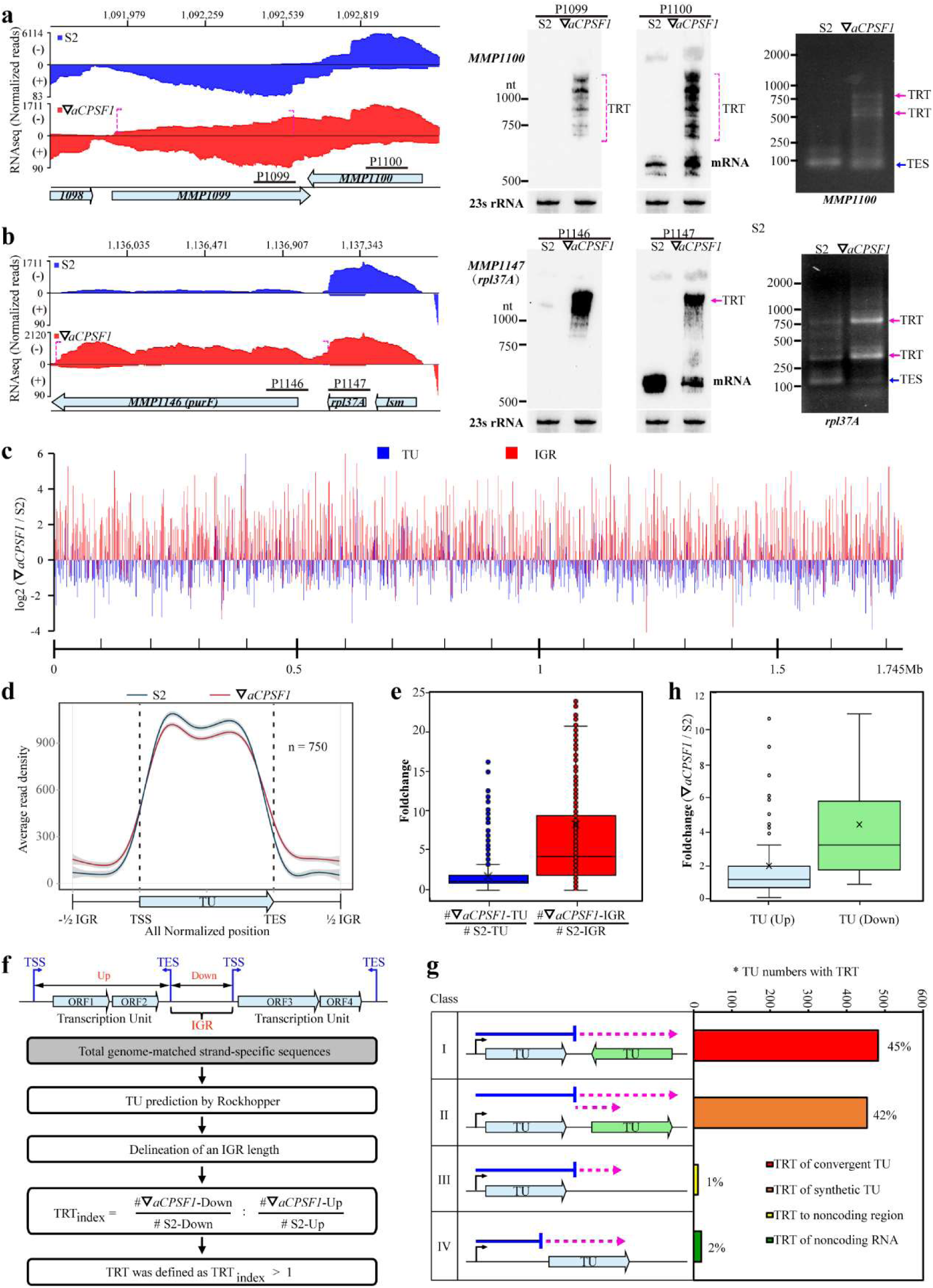
*Mmp-aCPSF1* depletion caused genome-wide transcription read-through (TRT). **a,b**, *Mmp-aCPSF1* depletion caused the 3′-end extensions (dotted magenta brackets) shown by strand-specific RNA-seq mapping profiles (left) of *MMP1100* (**a**) and *MMP1147* (**b**) in ▽*aCPSF1* (red) compared to that in strains S2 (blue). Numbers on the top indicate genomic sites, and bullets show genes. Northern blot (middle) assayed TRTs using the corresponding probes (horizontal sticks in **a**), and 23S rRNA was used as an internal control. 3′RACE (right) assayed the transcription end sites (TESs) and TRT extended regions. **c**, Pair-wise comparisons of the genome-wide Log2 FPKM ratios of transcription units (TUs) and intergenic regions (IGRs) between strains S2 and ▽*aCPSF1*. The ruler beneath indicates the genomic location. **d**, A metaplot diagram showing the average read mapping pattern of TUs and the flanking IGRs in strains S2 and ▽*aCPSF1* based on reads normalization of 750 TUs with an IGR length >100 nt. **e**, Boxplots showing the FPKM fold changes of 1,079 TUs and the flanking IGRs in ▽*aCPSF1* compared to that of S2, respectively. **f**, A flowchart depicting the procedure of TRT identification and TRT_index_ calculation. **g**, A diagram depicting the TRT types and percentages of each type occurred in ▽*aCPSF1*. Bent arrows and horizontal blue lines indicate TSS and transcript lengths in strain S2, respectively, and dot magenta arrows indicate transcripts that occur TRT in ▽*aCPSF1*. *, TU numbers that occur TRT. **h**, Boxplots showing the FPKM fold changes of Type II TRT in (**g**) for upstream TUs that generate TRTs (TU (Up)) and the tandem downstream TUs (TU (Down)), respectively.

### TRTs pervasively upregulate the expression of synthetic downstream transcripts and results in exuberant archaella along with enhanced motility

A pipeline (Fig. 2f) was developed to define *Mmp*-*aCPSF1* depletion caused TRT using four criteria: 1) 1,079 TUs were predicted based on transcriptome data; 2) delineation of IGR from upstream transcription end site (TES) and downstream transcription start site (TSS) on same DNA strand; 3) calculation of TRT_index_ by

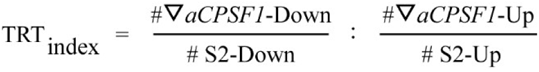

where, *#*▽*aCPSF1-* and #S2-Down indicate the IGR FPKMs in ▽*aCPSF1* and S2, respectively; while #▽*aCPSF1*- and #S2-Up refer to the TU FPKMs in ▽*aCPSF1* and S2, respectively; 4) TRT was defined as TRT_index_>1. Accordingly, TRT was identified in 965 TUs (89.5%) in ▽*aCPSF1*, of which 706 TUs (65.7%) have a TRT_index_ >2. Based on TRT location and direction, four types of TRT were classified as follows. TRT of upstream TU extended into the downstream oppositely encoded TU so to produce its antisense transcripts (type I, 45%), or into the downstream synthetic TU (type II, 42%), or just into the downstream IGR (type III, 1%); TRT of a IGR noncoding RNA extended into downstream TU (type IV, 2%) (Fig. 2g). Noticeably, type II TRTs markedly increased the transcriptions (median: 3.4-fold) of the tandem downstream TUs in ▽*aCPSF1* (Supplementary Dataset 2), compared to that of upstream TUs that generated TRTs (median: 1.3-fold) (Fig. 2h). Thus, type II TRTs severely disturbed downstream gene expression, and likely brought physiological changes.

The physiological change brought by type II TRTs was exemplified as that *MMP1719* TRT markedly increased transcription of the downstream archaella master regulator, *earA* (*MMP1718*) (Fig. 3 and Supplementary Dataset 1). Using three probes (P1, P2, P3) complementary to *MMP1717*, *earA*, and *MMP1719* respectively, Northern blot detected a long transcript encompassing *MMP1717*-*earA*-*MMP1719* in ▽*aCPSF1,* but only a co-transcribed TU of *MMP1717*-*earA* in S2 (Fig. 3b). 3′RACE also verified the 3′-extension of *MMP1719* TRT into *earA* and *MMP1717* (Fig. 3c). Moreover, a longer mRNA half-life (53.3 min) of the long *MMP1717*-*earA*-*MMP1719* transcript in ▽*aCPSF1* than the short *MMP1717*-*earA* transcript (11.2 min) in S2 was observed (Fig. 3d). Thereby, the markedly increased *earA* in ▽*aCPSF1* is likely derived from an overlay effect of *MMP1719* TRT and elevated stability of TRT generated transcript. Consistently, the *earA* targeted regulon^24^, the *fla* operon genes, were markedly upregulated in ▽*aCPSF1* (Fig. 3e and Supplementary Dataset 1). Northern blot detected the significant transcriptional increase of *flaB2* in ▽*aCPSF1*, but not in the double mutant of both *Mmp-aCPSF1* depletion and *earA* deletion (Fig. 3f), verifying that the increased *fla* operon expression in ▽*aCPSF1* was based on the upregulated expression of *earA*. Correspondingly, exuberant archaella and more active motility were observed in ▽*aCPSF1* cells compared with that in S2 (Fig. 3g, h). This indicated that *Mmp-aCPSF1* mediated transcription termination guarantees the optimal physiological process of archaea.

**Figure 3.**
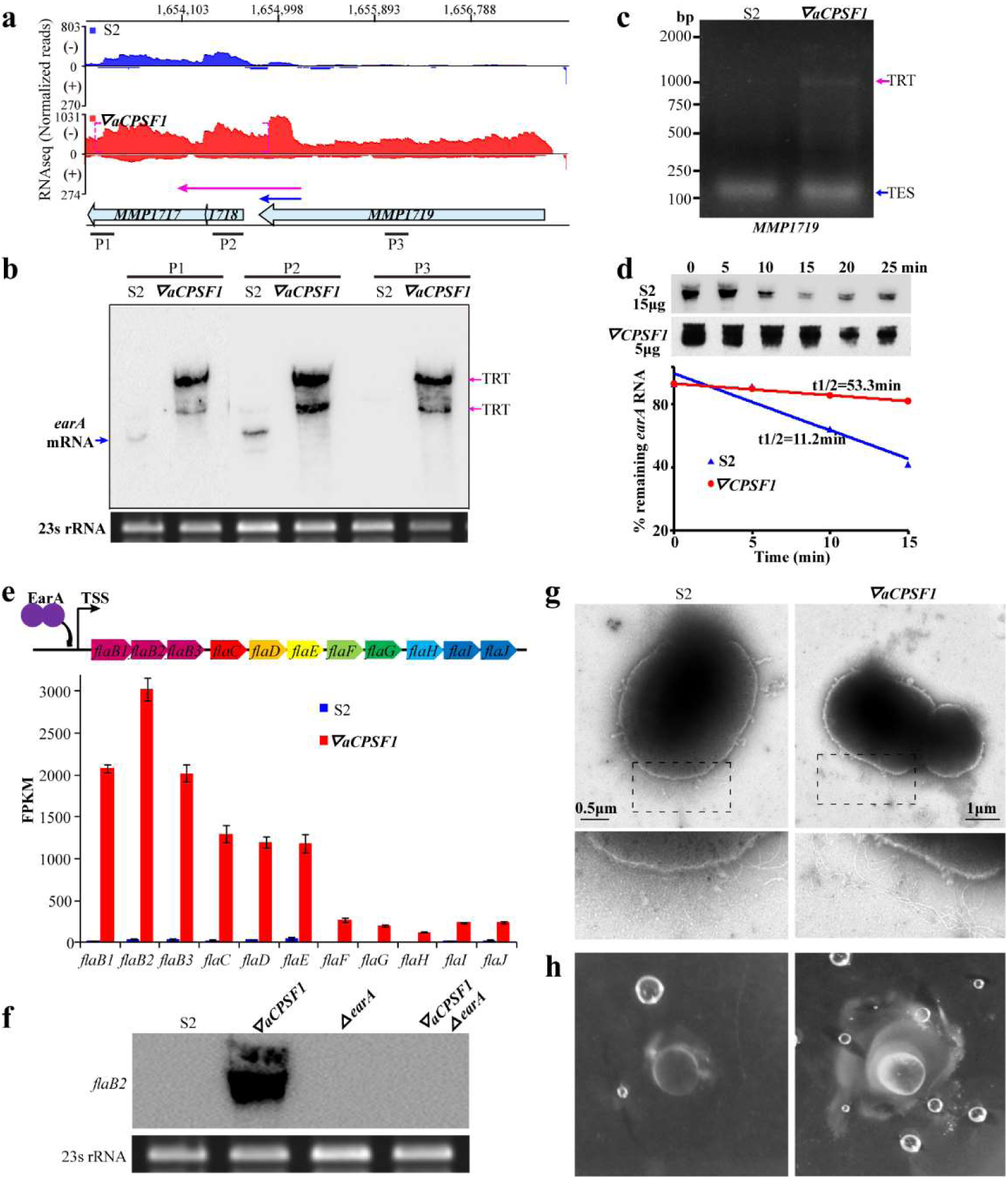
TRT upregulated expression of the archaella regulator *earA* resulted in exuberant archaella along with enhanced motility in ▽*aCPSF1*. **a**, RNA-seq reads mapped to *earA* (*MMP1718*) and the adjacent genes in strains S2 (blue) and ▽*aCPSF1* (red). The dot magenta bracket shows the 3′-end extension due to *Mmp-aCPSF1* depletion. Numbers on the top indicate the nucleotide sites of mapped genomic regions, and bullets represent genes. **b**, Northern blot assayed the *MMP1719* TRT extended into *earA* and *MMP1717* using three DNA probes P1, P2 and P3 which target the indicated genes. 23S rRNA is used as the internal control. **c**, 3′RACE amplified products containing normal transcription end site (TES) in S2 and TRT of *MMP1719* in ▽*aCPSF1*, respectively. **d**, Northern blot assayed *earA* transcript stability in strains S2 and ▽*aCPSF1* (upper panel). Half-lives (t1/2) are calculated based on the regression curve of remnant transcript content (lower panel). Loaded RNA contents of the two strains are indicated. Experiments were performed on triplicated cultures, and the average half-lives were shown. **e**, Upregulated transcription of the *fla* operon in ▽*aCPSF1*. A schematic depicts gene organization of the *fla* operon and its activator *earA* (upper) and the bar diagrams (lower) show the different FPKMs of each gene in strains S2 and ▽*aCPSF1*. **f**, Northern blot assayed the *flaB2* transcript in strains S2, ▽*aCPSF1*, the *earA* deletion (Δ*earA*), and a double mutation of *Mmp*-*aCPSF1* depletion and *earA* deletion (▽*aCPSF1*Δ*earA*). 23S rRNA was used as the sample control. **g**, Electron microscopic images display more archaella of ▽*aCPSF1* than that of strain S2 (upper) by amplified regions (dot framed) shown in lower panels. **h**, Mobility assay showed larger lawn and more bubbles in ▽*aCPSF1* grown 0.25% agar.

### Term-seq reveals a uridine-rich terminator motif and verifies the genome-wide TRT caused by *Mmp-aCPSF1* depletion

Next, we employed Term-seq^13, 25^ to sequence the transcript 3′-ends to determine the terminator characteristics of *M. maripaludis* and the function of *Mmp*-aCPSF1 in transcription termination. Based on stringent TES definition workflow described in Methods, 998 primary TESs were identified for 960 TUs and 38 non-coding RNAs (Supplementary Dataset 3), representing 67.7% (730) of the 1,079 predicted and 230 newly identified TUs in S2. Analysis of TES flanking sequences revealed a distinct upstream terminator motif embedding a 23 nt uridine-tract and the 4 nt most proximal TES having the highest match (Fig. 4a and Supplementary Dataset 3). By normalizing the read abundances of each 20 nt upstream and downstream of the 998 TESs, a >50% decrease between bases -2 and +2 flanking TES was found in S2 and defined transcription termination. However, <20% decrease at the two corresponding bases, and evidently higher reads were detected for the downstream 20 nts in ▽*aCPSF1* (Fig. 4b), manifesting a genome-wide TRT at single-base resolution. 3′RACE amplified the 3′-extensions for 19 TUs in ▽*aCPSF1* (Fig. 2a and Supplementary Fig. 2 and 4), and sequencing further validated the single-base accuracy and uridine-rich tracts of Term-seq defined TESs (Supplementary Fig. 5). These collectively indicate that the conserved archaeal ribonuclease, aCPSF1, is involved in genome-wide transcription termination.

**Figure 4.**
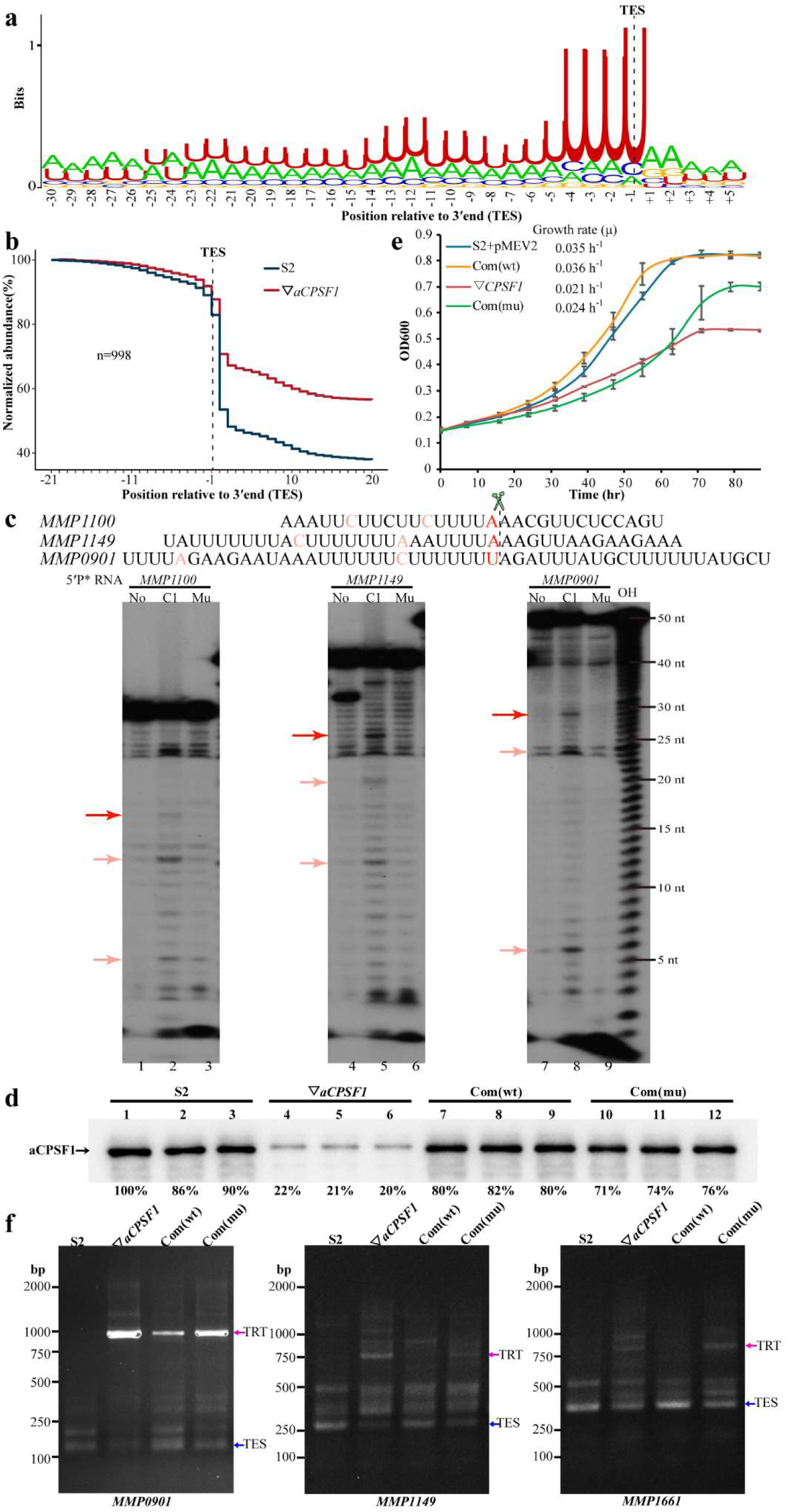
Terminator pattern of *M. maripludis* determined by Term-seq and evidence for *Mmp*-aCPSF1’s ribonuclease activity in transcription termination. **a**, The terminator signature of *M. maripaludis* S2 determined by Term-seq. A logo representation of the sequence motif upstream of 998 Term-seq defined TESs was generated using the MEME logarithm. **b**, A metaplot diagram showing the average reads of each 20 nt upstream and downstream of 998 primary TESs (dot black line) determined by Term-seq in strains S2 (blue line) and ▽*aCPSF1* (red line). **c**, The ribonuclease activity of *Mmp*-aCPSF1 on three representative RNA substrates derived from the sequences (upper) flanking the detected TESs (red bases) of *MMP1100*, *MMP1149,* and *MMP0901*. Urea sequencing gel displays the enzymatic cleavage results (lower). No, C1, and Mu indicate the nucleolytic assays without and with *Mmp*-aCPSF1 and its catalytic inactive mutant (H243A/H246A), respectively. Long and short red arrows indicate the major and minor cleavage products, respectively. OH, hydroxyl ladder that indicates RNA fragment migration positions. **d**, Western blot quantification of *Mmp*-aCPSF1 protein abundances in strains S2, ▽*aCPSF1*, *Mmp*-com(*Mmp*-C1) (Com(wt)), and *Mmp*-com(*Mmp*-C1mu) (Com(mu)). Triplicate results were showed. **e**, The growth curves of strains S2, ▽*aCPSF1*, Com(wt) and Com(mu) at 22°C. **f**, 3′RACE displaying the amplified TRTs in strains S2, ▽*aCPSF1*, Com(wt) and Com(mu). Blue and magenta arrows indicate the PCR products of normal terminations (TESs) and TRTs, respectively.

### *Mmp*-aCPSF1 depends on its ribonuclease activity to implement transcription termination

Nucleolytic activity of *Mmp*-aCPSF1 on TES motifs were then assayed using three representative uridine-rich RNA substrates (5′-[^32^P]-labeled), which were derived from the 3′-ends of *MMP1149* (50 nt), *MMP0901* (40 nt), and *MMP1100* (30 nt) (Fig. 4c upper). Addition of the overexpressed *Mmp*-aCPSF1, but not the activity-inactive mutant H243A/H246A (Supplementary Fig. 6), generated the major products with ∼25 nt, ∼20 nt, and ∼15 nt from the three synthetic RNAs, respectively, confirming that *Mmp*-aCPSF1 performs endoribonucleolytic activity at TESs downstream the uridine-tract terminators (Fig. 4c). Minor products cleaved after the upstream uridine-tract were also observed. Nevertheless, *Mmp*-aCPSF1 only exhibited weak ribonuclease activity *in vitro*, suggesting that interaction with other proteins or cofactors could promote its *in vivo* activity.

To evaluate the contribution of the ribonuclease activity of aCPSF1 in transcription termination *in vivo*, the wild-type *Mmp*-*aCPSF1* and its catalytic-inactive mutant, H243A/H246A, were expressed in ▽*aCPSF1* to construct two complementary strains, Com(wt): *Mmp*-com(*Mmp*-C1) and Com(mu): *Mmp*-com(*Mmp*-C1mu) (Supplementary Table 1) respectively. Although comparable cellular *Mmp*-aCPSF1 abundances were detected (Fig. 4d), Com(wt) completely restored the growth defect of ▽*aCPSF1* at 22°C, but Com(mu) exhibited similar reduced growth as ▽*aCPSF1* (Fig. 4e). Simultaneously, 3′RACE determined similar TRTs of *MMP0901*, *MMP1149*, and *MMP1661* in Com(mu) as in ▽*aCPSF1*, but not in Com(wt) (Fig. 4f). These suggested that *Mmp*-aCPSF1 implements transcription termination dependent on its ribonuclease activity.

### *Mmp*-aCPSF1 localizes within nucleoids and associates with archaeal RNA polymerase

A transcription termination factor must associate with RNAPs and generally also with chromosomal DNA inside the cell nuclei. Using cell-extract fractionation through size exclusion chromatography coupled with western blot, we detected the co-occurrence of *Mmp*-aCPSF1 and *Mmp*-RpoD, one of the 10 aRNAP subunits of *M. maripaludis,* in a protein complex of ∼200-660 kD. Ribonuclease digestion did not alter the co-occurrence (Fig. 5a), suggesting a nucleic-acid independent association of *Mmp*-aCPSF1 with *Mmp*-aRNAP. Using 3×Flag fused *Mmp*-aCPSF1 (*Mmp*-aCPSF1-F) and His_6_-3×Flag fused *Mmp*-RpoL (*Mmp*-RpoL-HF) strains (Supplementary Table 1), *Mmp*-aCPSF1 was immunoprecipitated from strain *Mmp*-aCPSF1-F and also detected in the His-affinity purification, co-immunoprecipitated, and tandem affinity chromatography products of strain *Mmp*-RpoL-HF (Fig. 5b), and RNase, DNase, and Benzonase nuclease (digesting both DNA and RNA) did not alter the results (Fig. 5c), thus verifying the *in vivo* direct association of *Mmp*-aCPSF1 and *Mmp*-aRNAP. Using super-resolution photoactivated localization microscopy (PALM) imaging, the mMaple3 fluorescence signal of the mMaple3 fused *Mmp*-RpoL (*Mmp*-RpoL-mMaple) was observed within the nucleoid throughout the growth, while that of *Mmp*-aCPSF1-mMaple was detected primarily (∼77%) in the nucleoid at the early-exponential cells (OD_600_ ∼ 0.2) (Fig. 5d and Supplementary Fig. 7). Thus, a direct association with aRNAP and the nucleoid localization both warrant *Mmp*-aCPSF1 to participate in transcription.

**Figure 5.**
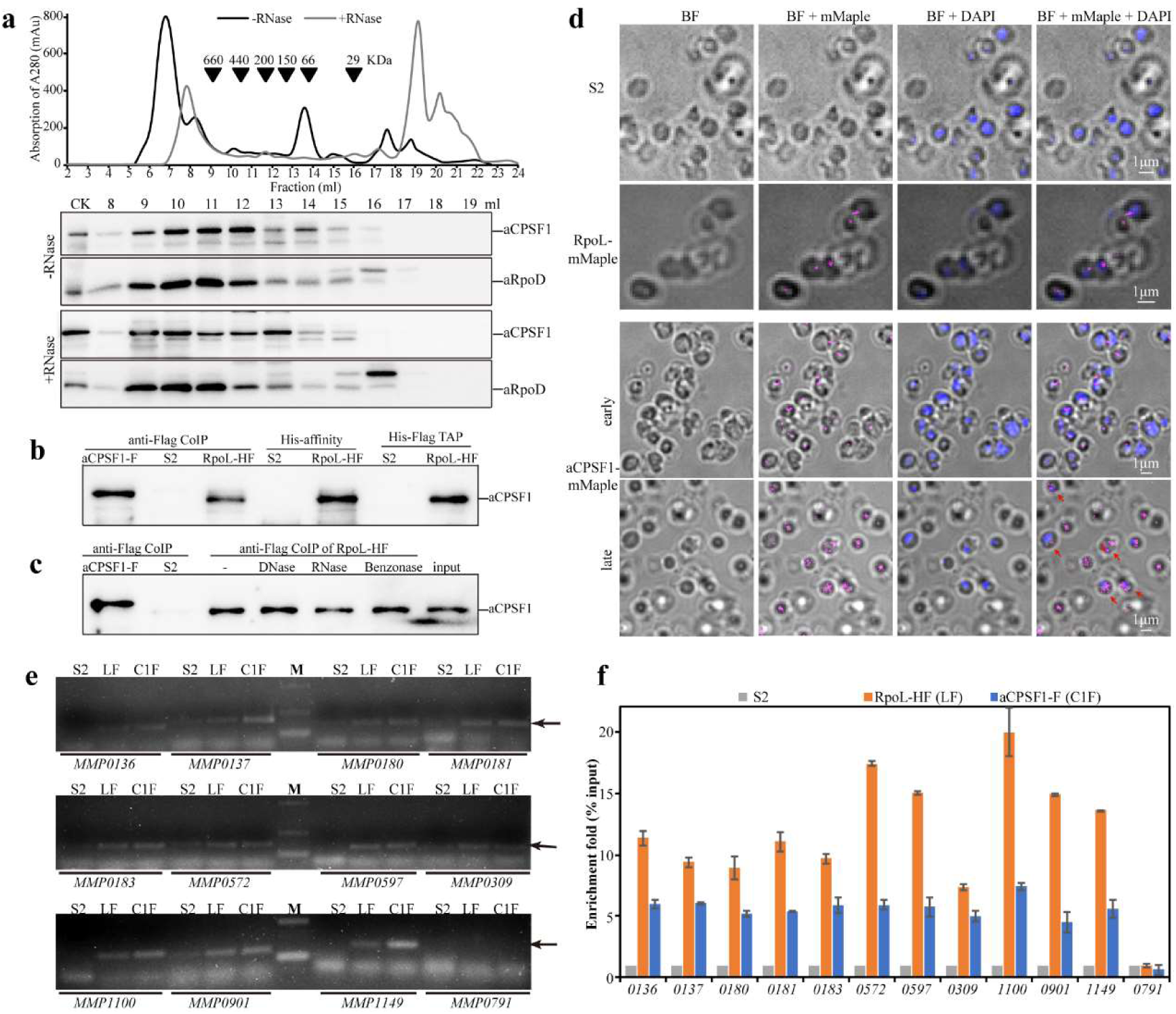
Association of *Mmp*-aCPSF1 with *Mmp*-aRNAP and its nucleoid and chromosomal localizations. **a**, Co-occurrence of *Mmp*-aCFSF1 and *Mmp*-RpoD assayed by *M. maripaludis* cell extract fractionation via size exclusion chromatography (upper) coupled with western blot analysis (lower). +/–RNase, with or without RNase A treatment. **b**,**c**, Association of *Mmp*-aCPSF1 with *Mmp*-aRNAP determined through co-immunoprecipitation (anti-Flag CoIP), Ni column affinity (His-affinity), or tandem affinity purification (TAP) using strains *Mmp*-RpoL-HF and *Mmp*-aCPSF1-F. Strain S2 was used as a mock control. Western blot determined the presence of *Mmp*-aCPSF1 in the captured products. -, RNase, DNase, and Benzonase indicate treatments without and with RNase A, DNase I, and Benzonase nuclease to the *Mmp*-RpoL-HF cell extract before immunoprecipitation, respectively. **d**, Cellular localization of mMaple3 fused *Mmp*-aCPSF1 (aCPSF1-mMaple) and *Mmp*-RpoL (RpoL-mMaple) at early- or late-exponential phases determined by super-resolution PALM imaging. Strain S2 was included as control. BF, bright field; BF+mMaple, overlay of bright field and mMaple3 images; BF+DAPI, overlay of bright field and DAPI images; BF+mMaple+DAPI, overlay of bright field, mMaple3 and DAPI images. Arrows indicate the mMaple3 signals in cytoplasm. **e**, PCR products (arrows pointed) of the indicated genes amplified from the ChIP DNAs from strains S2 (mock), *Mmp*-RpoL-HF (LF), and *Mmp*-aCPSF1-F (C1F), respectively. M, DNA marker. **f**, Enrichment ratio (% input) was calculated as the qPCR assayed gene copies in ChIP products over that in total DNA (input), and enrichment fold was defined as the enrichment ratios of *Mmp*-RpoL-HF and *Mmp*-aCPSF1-F to that of S2 (mock control).

### *Mmp*-aCPSF1 has a chromosomal occupancy at transcript 3′-ends

Using chromatin immunoprecipitation (ChIP), the chromosomal DNA occupancy of *Mmp*-aCPSF1 was determined. The chromatin DNAs captured by Flag-tagged *Mmp*-aCPSF1 or aRNAP were immunoprecipitated and used as PCR templates. Abundant PCR products in length 130–150 bp were amplified for 11 efficiently expressed TUs (Fig. 5e). qPCR verified ∼6- and 6–12-fold enrichments in the 3′-end regions for these TUs from the Flag-tagged *Mmp*-aCPSF1- and *Mmp*-RpoL-captured DNAs respectively, compared with that from a mock ChIP of strain S2 (Fig. 5f). No PCR products were amplified from the poorly expressed *MMP0791* (Fig. 5e, f). Collectively, *Mmp*-aCPSF1 associates with aRNAP as well as chromosomal DNA, the key features of a transcription termination factor.

### aCPSF1 appears to co-evolve with aRNAP and its targeted uridine-rich motif widely distributes among archaeal IGRs

To explore the universality of aCPSF1-mediated transcription termination among Archaea, aCPSF1 orthologs of 31 representative species from all four defined archaeal superphyla (Euryarchaeota, TACK, Asgard and DPANN) were subjected to phylogenetic analysis (Fig. 6a). Simultaneously, a phylogenetic analysis was conducted on a protein concatenation of aRNAP subunits A, B, and E, and the conserved transcription elongation factor Spt5. aCPSF1 exhibited as a good phylogenetic marker providing an equivalent phylogenetic topology as that of the concatenated sequences of 26 universally conserved proteins^26^, and also showed convergent phylogenetic clustering as aRNAPs (Fig. 6a). Thus, aCPSF1 and RNA transcription machinery are likely to have co-evolved.

**Figure 6.**
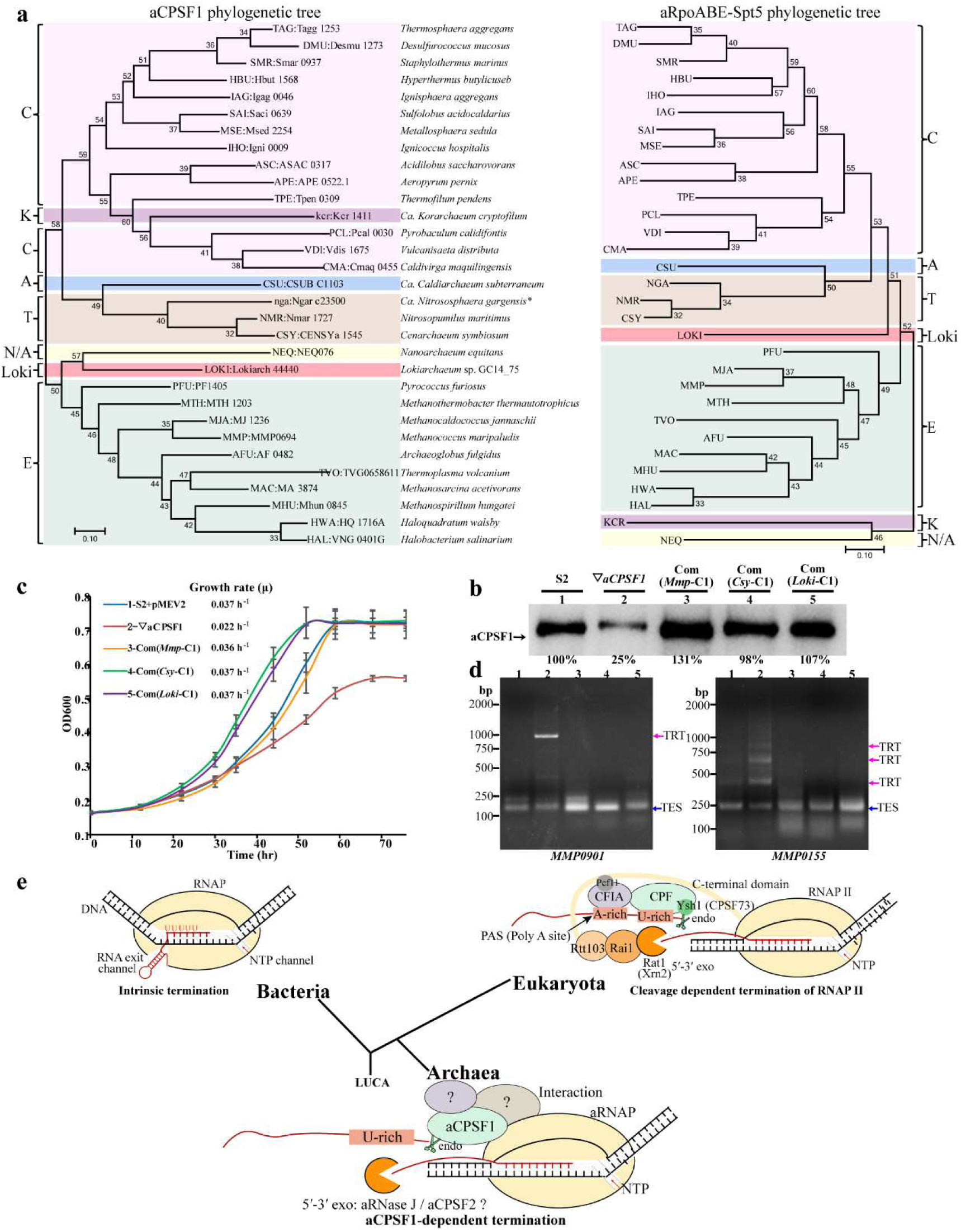
The conserved aCPSF1 coevolves with the transcription machinery and likely represents a primordial and simplified transcription termination of the Eukaryotic RNAP II mode. **a**, The Maximum Likelihood phylogenetic trees based on aCPSF1 proteins (left) and concatenated RNA polymerase subunits A, B and E, and transcription elongation factor Spt5 (right). Protein sequences were retrieved from the genomes or representative metagenome-assembled genomes of archaeal lineages that have been described thus far, including Euryarchaeota (E), Crenarchaeota (C), Thaumarchaeota (T), Aigarchaeota (A), Korarchaeota (K), Nanoarchaeota (N), and Lokiarchaeota (Loki). The unrooted tree was constructed using Maximum-Likelihood methods in MEGA 7.0 software package. Bootstrap values are shown at each node. b, Ectopic expression of the aCPSF1 orthologs from *Lokiarchaeum* GC14_75 (com(*Loki*-C1)) and *Cenarchaeum symbiosum* (com(*Csy*-C1)) restored the growth defects of the methanococcal ▽*aCPSF1* at 22°C to similar growth rates as that of the complementation of itself (com(*Mmp*-C1)) and strain S2 carrying an empty vector (S2-pMEV2). c, *Mmp*-aCPSF1 protein contents in various ortholog complementary strains examined by western blot analysis using its polyclonal antiserum. Percentiles of aCPSF1 protein content in reference to that in S2 are shown beneath the gel. d, 3′RACE assays of TRT transcripts of *MMP0901* and *MMP0155* in the indicated strains by the numbers as in panel b. e, A proposed aCPSF1 dependent transcription termination mechanism in Archaea that features the evolutionary relics of the bacterial intrinsic^2, 3^ and eukaryotic RNAP II termination mechanisms^3–5^. LUCA, last universal common ancestor.

Given that aCPSF1 acts on uridine-rich terminator motif, distributions of the uridine-rich sequences were searched at the 200 nt upstream and downstream the predicted stop codons of annotated ORFs in the 31 representative archaeal species. Notably, the result showed that compared with the upstream regions, the uridine-rich tracts (U5, U6 and U7) were significantly and specifically enriched in the IGRs in all detected archaeal species (Supplementary Fig. 8 and Dataset 4). This indicates that prevalent uridine-rich tract in IGRs could be a conserved terminator sequence of archaea. The high conservation of both aCPSF1 and the uridine-rich motif supports that aCPSF1 dependent transcription termination could be universally employed in Archaea.

### Two phylogenetic distant *aCPSF1* orthologs function in methanococcal transcription termination

To experimentally verify the conserved function of aCPSF1 in transcription termination, two *aCPSF1* orthologs respectively from Ca. *Lokiarchaeum* sp. GC14_75 (Lokiarchaeota), and Ca. *Cenarchaeum symbiosum* (Thaumarchaeota) were codon optimized (Supplementary Fig. 9) and ectopically expressed in ▽*aCPSF1* (Fig. 6b). Behaving as *Mmp*-aCPSF1, they not only restored the cold-sensitive growth defects of ▽*aCPSF1* (Fig. 6c), but also eliminated the TRTs of convergent (*MMP0901*, type I) and synthetic TUs (*MMP0155*, type II) (Fig. 6d). This suggested that aCPSF1s from various archaeal phyla play a conserved role in transcription termination.

## Discussion

Up to date, transcription termination mechanisms remain almost unknown and no general transcription termination factor was reported for Archaea^1, 13, 18^. In this work, through extensive genetic, molecular and biochemical experiments, and high-throughput RNA-seq and Term-seq, we have revealed that the archaeal conserved ribonuclease aCPSF1, by cleaving on the transcript 3′-end uridine-rich sequences (Fig. 4), controls the genome-wide transcription termination (Fig. 2) and ensures the programmed transcriptome and so the optimal physiology and growth (Fig. 1 and 3) in a methanoarchaeon. The nucleoid localization and direct association with archaeal RNAP as well as the chromosome DNA of aCPSF1 (Fig. 5) all ensure its direct action on transcription termination. Thus, aCPSF1 is the first reported general transcription termination factor of Archaea, and the essential role in transcription termination discovered here deciphers its universality and essentiality in Archaea.

### *Mmp*-aCPSF1-mediated transcription termination is essential for archaeal programmed transcriptome

Due to the short IGRs within the prokaryotic compact genomes, transcription termination for ordered transcriptomes should be more vital. This is verified by that *Mmp*-*aCPSF1* depletion caused genome-wide TRTs markedly perturb the ordered transcriptome in particularly increasing the transcription of downstream genes (type II TRT, 42%) (Fig. 1 and 2). This concomitantly changed the normal physiology exemplified as exuberant archaella along with promoted motility caused by TRT increased expression of the archaella master regulator, *earA* (Fig. 3). Moreover, *Mmp-aCPSF1* depletion inversely changed the expression of the highly- and poorly-transcribed genes grouped in wild-type behaving as a “robbing the rich to feed the poor” pattern (Fig. 1e, f), resulting an overall transcriptome chaos and a concomitant cold-sensitive growth (Fig. 1). We postulated this is because that *Mmp*-*aCPSF1* depletion caused abnormal termination could sequester RNAP for recycling for new rounds of transcription. These results highlight the significance of aCPSF1 dependent transcription termination in maintaining the ordered transcriptome and optimal physiology/growth of Archaea. Similar outcomes of transcription termination defects are also found in Bacteria^27, 28^ and even Eukaryotes^8, 29, 30^.

### Both uridine-tract terminator signal and aCPSF1 seem to be required for archaeal transcription termination

Term-seq analysis determines a two-consecutive uridine-tracts terminator motif in *M. maripaludis* (Fig. 4a). Such similar but slight distinct motifs were also observed in two represent Euryarchaeota and Crenarchaeota species, *Methanosarcina mazei* and *Sulfolobus acidocaldarius*^13^. Uridine-rich sequences were considered as the intrinsic termination signal based on the earlier and mainly *in vitro* studies^16, 17, 31^. However, whether the uridine-rich sequence is merely to pause aRNAP or also dismantle it to dissociation and termination has not been clarified. In this work, *Mmp*-aCPSF1 was demonstrated to perform endoribonucleolytic cleavage downstream the uridine-rich sequences to produce putative TESs *in vitro*, and this endoribonuclease activity was determined essential for transcription termination *in vivo* as well (Fig. 4). Thus, both uridine-rich terminator signal and aCPSF1 seem to be required for aRNAP transcription termination, and based on these results, a model was proposed (Fig. 6e). The intrinsic uridine-rich terminator signal could cause the progressing aRNAP to pause and buy time for aCPSF1 to recognize and cleave downstream it, and trigger aRNAP to dissociation and so transcription termination. The nucleoid location and direct association of aCPSF1 with aRNAP (Fig. 5a-d) would guarantee the recruitment of aCPSF1 to transcription machinery and uridine-rich signal on nascent RNAs. aCPSF1 exhibits as a conserved phylogenetic marker and convergent phylogenetic clustering as aRNAP (Fig. 6a), in addition that uridine-rich motif was found prevalent in IGRs among representative archaeal genomes (Supplementary Fig. 9 and dataset 4). Moreover, two distant aCPSF1 orthologs each from Lokiarchaeota and Thaumarchaeota could implement transcription termination in *M. maripaludis* (Fig. 6b-d). Consequently, we predict that the aCPSF1 mediated transcription termination could emerge predating the divergence of various archaeal phyla and is still wildly employed in modern Archaea.

### The archaeal 3′-end cleavage dependent transcription termination could expose an archetype of eukaryotic RNAP II termination mode

Strikingly, the working mechanism of aCPSF1 that cleaves at transcript 3′-ends to terminate transcription resembles the eukaryotic 3′-end processing/cleavage triggered RNAP II termination, wherein the aCPSF1 homologs, the yeast Ysh1 and the human CPSF73, in the multi-subunits 3′-end processing machinery execute the 3′-end cleavage^3, 5, 8, 32, 33^. The uridine-rich sequences that are recognized by the accessory cleavage factors IA and IB (CFIA and CFIB) in 3′-end processing machinery are also determined necessary to facilitate Ysh1 for 3′-end cleavage in yeast^32^. While aCPSF1 could probably rely on its N-terminal two K homology (KH) domains^20, 21^ to recognize the uridine-rich signal by itself. Interestingly, one supposed archaeal ancestor of eukaryotes^34^, Lokiarchaeota, appears to use the aCPSF1 termination mechanism as well, which is evidenced by *Locki*-aCPSF1 terminating the methanococcal transcription and also possessing the genome-wide prevalent IGR uridine-rich sequences. Therefore, it is assumed that the eukaryotic RNAP II termination could be originated from the aCPSF1 dependent mode reported here and this could be one more evidence for that Archaea is the latest ancestor of eukaryotes. Distinctly, CPSF73 cleaving downstream of poly(A) site of transcript 3′-end not only terminates transcription but also provides a polyadenylation site for transcript maturation^33, 35^, while no 3′ terminus polyadenylation followed aCPSF1 3′-end cleavage. Moreover, very few homologs of the multi-subunit 3′-end processing complex^36, 37^ and the associated eukaryotic termination factors^8–11^ were identified in Archaea. The 5′-3′ eukaryotic exoribonuclease, yeast Rat1 or human Xrn2, has been determined to be recruited to the 5′-end of the cleaved downstream RNA that are still loaded by RNAP, and rapidly ‘chew’ its way along the RNA to catch up and dismantle the preceding RNAP, therefore working as a “torpedo” for eukaryotic RNAP dissociation and termination ^8, 9, 12, 29^. Although no homologs of Rat1 or Xrn2 were found in Archaea, 5′-3′ exoribonucleases aRNase J, aCPSF2 and even aCPSF1 itself were indeed identified ^19, 20, 38, 39^. They may play comparable function in aRNAP dissociation and termination in Archaea (Figure 6e). Therefore, aCPSF1-mediated archaeal transcription termination could employ other or none protein factors, so would represent a simplified mode and an evolutionary relic of eukaryotic RNAP II termination mechanism. Additionally, the uridine-rich tracts at proximal transcript 3′-ends that are recognized by aCPSF1 in Archaea appears to have been adopted as an element of bacterial intrinsic terminator (Fig. 6e), which represents shared feature of the bacterial and archaeal terminator motifs. Thus, the terminator uridine-rich tract and aCPSF1 cleavage dependent termination mode of Archaea could reveal a genetic jigsaw of the bacterial and eukaryotic termination mechanisms; while the 3′ transcript cleavage but no polyadenylation could expose a primordial termination strategy predating the divergent evolution of Archaea and Eukaryotes. Therefore, insight into aCPSF1 mediated archaeal transcription termination should shed lights on understanding both the complex eukaryotic transcription termination mechanism and the evolutionary trajectory of life’s transcription process.

### The involvement of aCPSF1 in RNA decay

Moreover, *Mmp*-aCPSF1 also appears to function in RNA degradation and turnover, as its depletion also results in approximate 2-fold prolonged bulk mRNA lifespans at 22°C (Fig. 1c). The cytoplasmic location of *Mmp*-aCPSF1 during later growth phases (Fig. 5d and Supplementary Fig. 7) and the intrinsic nature of endoribonuclease and exoribonuclease activities ^19–21^ could support its role in regular RNA turnover. As no exosome subunits were identified in methanococcal species ^40^, aCPSF1 as well as other ribonucleases of β-CASP family were supposed to play major roles in RNA turnover ^39, 41, 42^. While this work provides evidence for the involvement of aCPSF1 in RNA degradation. The role of aCPSF1 involved in RNA decay, another one of life’s fundamental biological processes, deciphers its universality and essentiality in all archaeal phyla as well.

## Methods

### Strains, plasmids and culture conditions

Strains used in this study are listed in Supplementary Table 1. *M. maripaludis* S2 and its derivatives were grown in pre-reduced McF medium under a gas phase of N2/CO2 (80:20) at 37°C and 22°C as previously described^43^, and 1.5% agar was used in solid medium. Puromycin (2.5 μg/ml), neomycin (1.0 mg/ml) and 8-azahypoxanthine (0.25 mg/ml) were used for genetic modification selections unless indicated otherwise. *E. coli* DH5α, BL21(DE3)pLysS and BW25113 were grown at 37°C in Luria-Bertani (LB) broth and supplemented with ampicillin (100 μg/ml) or streptomycin (50 μg/ml) when needed.

Construction of the *Mmp-aCPSF1* gene depletion and complementary strains.

Given the possibly essentiality of *Mmp-aCPSF1* determined in *M. maripaludis*^22^ and repeated failures to delete *Mmp-aCPSF1* from the chromosome, we constructed the *Mmp-aCPSF1* (*MMP0694*) expression depleted strain (▽*aCPSF1*) using a *TetR*-*tetO* repressor-operator system as follows.

First, a tetracycline responsive regulator (*tetR*) and operator (*tetO*) cassette from *Bacillus subtilis* was fused upstream to the N-terminal His6-tagged *Mmp*-*aCPSF1*, and then transformed and integrated at the *hpt* locus of *M. maripaludis* S2 to construct a tetracycline regulated *Mmp*-*aCPSF1* overexpression strain *tetO-aCPSF1* by 8-azahypoxanthine sensitivity selection. To achieve gene expression in *M. maripaludis*, the promoter (P*mcr*) and terminator (T*mcr*) of the methyl-CoM reductase were fused to the *tetR* gene at the 5′- and 3′-termini, respectively, and inserted to pMD19-T (TAKARA) to obtain pMD19-T-P*mcr*-*tetR*-T*mcr* (Supplementary Table 1). The *tetO* operator was positioned downstream the nitrogen fixation gene *nif* promoter (P*nif*) and fused upstream the amplified *Mmp-aCPSF1* ORF to obtain a *tetO* fused *Mmp-aCPSF1* fragment P*nif*-*tetO*-His6-*aCPSF1*. For homologous recombination at *hpt* site, each of approximate 800 bp fragment upstream and downstream *hpt* was amplified from the S2 genomic DNA and fused with the P*mcr*-*tetR*-T*mcr* and P*nif*-*tetO*-His6-*aCPSF1* fragment at each end to obtain the plasmid of p*tetR*/*tetO*-His6-*aCPSF1* (Supplementary Table 1) via stepwise Gibson assembly using ClonExpress MultiS One Step Cloning Kit (Vazyme). The PCR fragment amplified from the plasmid p*tetR*/*tetO*-His6-*aCPSF1* using primers *Hpt*up-F/*Hpt*down-R (Supplementary Table 2) was transformed into S2 using polyethylene glycol (PEG)-mediated transformation approach ^44^ to generate a tetracycline regulated *Mmp*-*aCPSF1* overexpression strain (*tetO-aCPSF1*) via double cross homologous recombination. Transformants were selected by plating on McF agar containing 8-azahypoxanthine as described previously ^45^, and confirmed via PCR amplification using primers *Hpt*up-F and *Hpt*down-R.

Next, an in-frame deletion of the indigenous *Mmp*-*aCPSF1* gene was implemented in strain *tetO-aCPSF1*. An approximate 800 bp DNA fragment each upstream and downstream the *Mmp*-*aCPSF1* gene was PCR amplified and fused to the puromycin-resistance cassette *pac* gene amplified from plasmid pIJA03 at each end to generate plasmid pMD19T-Δ*aCPSF1* via stepwise Gibson assembly. The DNA fragment of *aCPSF1*up-*pac*-*aCPSF1*down amplified from pMD19T-Δ*aCPSF1* was transformed into strain *tetO-aCPSF1* to knockout the indigenous *Mmp*-*aCPSF1*. Transformants were selected on McF agar containing puromycin, and the *Mmp-aCPSF1* gene deletion was validated by PCR amplification. Thus, a tetracycline regulated *Mmp-aCPSF1* expression strain ▽*aCPSF1* was obtained.

To obtain various ▽*aCPSF1* complementary strains, corresponding *aCPSF1* genes were cloned into the *Xho*I/*Bgl* II sites of expression plasmid pMEV2 to produce: pMEV2-*aCPSF1*(wt) / pMEV2-*aCPSF1*(mu) carrying the wild-type *Mmp-aCPSF1* and its mutant, H243A/H246A, respectively, and pMEV2-*Lokiarch44440* / pMEV2*-CENSYa1545*, carrying two codon optimized *Mmp-aCPSF1* orthologs from Ca. *Lokiarchaeum* sp. GC14_75 and Ca. *Cenarchaeum symbiosum*. The constructed plasmids were each transformed into ▽*aCPSF1* via PEG-mediated transformation approach to produce the complementary strains.

### Construction of the in-frame deletion mutants of *earA* in strains S2 and ▽*aCPSF1*

The in-frame deletion mutants of *earA* were constructed similarly as above. Each of approximate 800 bp DNA fragment upstream and downstream *earA* (*MMP1718*) was amplified and fused to a neomycin-resistance cassette amplified from the plasmid pMEV2. The resulted *earA*up-*neo*-*earA*down fragment was transformed into strains S2 and ▽*aCPSF1*, and via double cross homologous recombination, the *earA* deletion mutants Δ*earA* and ▽*aCPSF1*Δ*earA* (Supplementary Table 1) were obtained, respectively.

### Construction of Flag-tagged *Mmp-*aCPSF1 and His-Flag-tagged *Mmp-*RpoL and mMaple3 fusion strains

3×Flag or His6 and 3×Flag tags were firstly fused to the C termini of *Mmp*-aCPSF1 and *Mmp*-RpoL via PCR amplification and cloned into pMD19-T to construct the plasmids pMD19-T-*aCPSF1*-3Flag and pMD19-T-*RpoL*-3Flag, respectively. The puromycin or neomycin cassette in plasmid pIJA03 or pMEV2 was amplified and overlapping fused to the ∼800bp downstream of *Mmp-aCPSF1* or *Mmp-RpoL*. The overlapped PCR fragments were then inserted into downstream of *Mmp*-*aCPSF1*-3Flag or *Mmp*-*RpoL*-3Flag to obtain pMD19-T-*aCPSF1*-3Flag-*pac* and pMD19-T-*rpoL*-3Flag-His_6_-*neo* (Supplementary Table 1), respectively. Next, DNA fragments were amplified using primers *aCPSF1*Flag-F/*aCPSF1*Flagdw-R and *RpoL*Flag-F/*RpoL*Flagdw-R from pMD19-T-*aCPSF1*-3Flag-*pac* and pMD19-T-*rpoL*-3Flag-His_6_-*neo*, respectively, and transformed into *M. maripaludis* S2 to obtain Flag-tagged strains of *Mmp*-aCPSF1-F and *Mmp*-RpoL-HF. The chromosomal mMaple3 fusions at the C-termini of *Mmp*-RpoL and *Mmp*-aCPSF1 were constructed similarly except for using mMaple3 sequence to replace that of Flag-tag and resulted in strains of *Mmp*-aCPSF1-mMaple and *Mmp*-RpoL-mMaple (Supplementary Table 1), respectively.

### RNA-seq and data analysis

Using the methods as described previously^46, 47^, high-throughput strand-specific RNA-seq were performed for differential transcriptome analysis between strains S2 and ▽*aCPSF1* on biological triplicates. The uniquely mapped reads were aligned to *M. maripaludis* S2 reference genome (GCF_000011585). Differential expression analysis of strains S2 and ▽*aCPSF1* was performed using the DESeq algorithm with the normalized mapped read counts, and genes with an adjusted P-value < 0.05 in DESeq were assigned as differentially expressed^48, 49^.

To calculate the genome-wide Transcriptional Read-through index (TRT_index_), transcriptional units (TUs) were first predicted by Rockhopper. Intergenic regions (IGRs) were assigned as from the transcription end sites (TESs) to the transcription start sites (TSSs) of downstream TUs on same strand. Transcript abundances of each TU and IGR in strains S2 and ▽*aCPSF1* were quantified using stringtie (v1.3.3) with –e option, and transformed to GTF file format used as –G parameter, and normalized by total sequence reads depth determined by samtools (v1.3.1). Differential expression analysis of TUs and IGRs in strains S2 and ▽*aCPSF1* was performed using the DESeq R package (1.18.0).

### Term-seq and data analysis

Term-seq libraries were constructed by following the procedure described previously^25^. Sequencing was performed on an Illumina HiSeq X-ten system with 150-bp paired-end reads. After quality control processing of raw data, filtered reads were aligned to the *M. maripaludis* S2 reference genome (GCF_000011585) using Bowtie^50^. The read accounts of the 3′-ends mapped to each genomic position in strain S2 were recorded. TESs of each gene were assigned based on four criteria: 1). within 200 nt of the downstream region distant to the stop codon of a gene; 2) >1.1 reads ratio of -1 site (upstream TES) to +1 site (predicted TES); 3) read-counts of -1 site minus +1 site > 5; 4) upon 2) and 3) satisfied, the site that has the highest readcounts difference of -1 site minus +1 site was recorded as a primary TES (Supplementary Database 3). Using MEME algorithm^51^, terminator motifs were searched in the sequences flanking Term-seq predicted TES and sequence logos were generated by WebLogo^52^. To compare the differences of reads coverage flanking the primary TESs in strains S2 and ▽*aCPSF1*, a metaplot analysis was performed on the average reads of 20 nt each upstream and downstream of all the primary TESs and normalized by dividing that of the first site of 20 nt upstream TESs.

### Rapid amplification of cDNA 3′-ends (3′RACE)

3′RACE was performed as described previously^53^. 20 μg of total RNA were ligated with 50 pmol 3′adaptor Linker (5′-rAppCTGTAGGCACCATCAAT–NH2-3′; NEB) through a 16 h incubation at 16°C with 20 U T4 RNA ligase (Ambion). 3′ linker-ligated RNA was recovered by isopropanol precipitation and one aliquot of 2 μg was mixed with 100 pmol 3′R-RT-P (5′-ATTGATGGTGCCTACAG-3′, complementary to the 3′ RNA linker) by incubation at 65°C for 10 min and on ice for 2 min, and was used in reverse transcription (RT) reaction using 200 U of SuperScript III reverse transcriptase (Invitrogen). After RT, nested PCR was conducted using primers (Supplementary Table 2) targeted regions of <200 nt upstream the termination codons to obtain gene-specific products. Specific PCR products were excised from 2% agarose gel and cloned into pMD19-T (TaKaRa) and then sequenced. The 3′-end of a TU is defined as the nucleotide linked to the 3′RACE-linker.

### Protein purification

For purification of *Mmp*-aCPSF1 protein, a His-tagged SUMO-*aCPSF1* fusion was first constructed and cloned into pSB1s using the ClonExpress MultiS One Step Cloning Kit. Plasmid pSB1s carrying His-SUMO-*aCPSF1* was then transformed into *E. coli* BW25113 and the transformants were induced by 0.1% L-arabinose. After a 16 h-induction at 22°C, cells were harvested, and the protein was purified as described previously^39^. Next, His-SUMO tags were removed by incubated with His-tagged Ulp1 protease at 30°C for 2h, and *Mmp*-aCPSF1 protein was purified through a His-Trap HP column to remove His-tagged SUMO and His-tagged Ulp1. The Rpo D/L protein of *M. maripaludis* S2 (*Mmp*-aRpoD/L) was overexpressed and purified as described previously^54^. In brief, the ORFs of *Mmp*-*rpoD* (*MMP1322*) and *Mmp*-*rpoL* were stepwise cloned into pGEX-4T-1 to get the plasmid pGEX-4T-1-*rpoD/L*, and transformed into *E. coli* BL21(DE3)pLysS for overexpression. Purified proteins were detected by SDS-PAGE, and protein concentration was determined using a BCA protein assay kit (Thermo Scientific).

### Nuclease activity assay for *Mmp*-aCPSF1

Three RNA fragments containing the termination sites and flanking sequences of *MMP1100, MMP1149* and *MMP0901* (Fig. 4c) were synthesized by Genscript (China). They were 5′-end labeled by [λ-^32^P] ATP (PerkinElmer) using T4 polynucleotide kinase (Thermo Scientific). A 10 μl nuclease reaction mixture contained 20 μM wild-type *Mmp*-aCPSF1 protein or its catalytic inactive mutant, 5 nM 5′-[γ^32^P] labeled RNA, 20 mM HEPES, (pH 7.5), 150 mM NaCl, 5 mM MgCl_2_ and 5% (w/v) glycerol. Nucleolytic reactions were initiated by the addition of the protein at 37°C for 90 min, and stopped by incubation with 10 μg/ml Proteinase K (Ambion) at 55°C for 15 min. The reaction products were then mixed with formamide-containing dye and analyzed on 10% sequencing urea-PAGE. A nucleotide ladder was generated by alkaline hydrolysis of the labeled RNA substrates. The urea-PAGE gels were analyzed by autoradiography with X-ray film.

### Western blot assay

Western blot was performed to determine the cellular *Mmp*-aCPSF1 or *Mmp*-Rpo D/L protein abundances in various genetic modified strains. A polyclonal rabbit antiserum against the purified *Mmp*-aCPSF1 or *Mmp*-Rpo D/L protein was raised by MBL International Corporation, respectively. The mid-exponential cells of *M. maripaludis* were harvested and resuspended in a lysis buffer [50 mM Tris-HCl (pH 7.5), 150 mM NaCl, 10 (w/v) glycerol, 0.05% (v/v) NP-40], and lysed by sonication. Cell lysate was centrifuged and the proteins in supernatant were separated on 12% SDS-PAGE and then transferred to nitrocellulose membrane. The antisera of anti-*Mmp*-aCPSF1 (1: 8,000) or anti-*Mmp*-aRpoD/L (1: 20000) were diluted and used respectively, and a horseradish peroxidase (HRP)-linked secondary conjugate at 1: 5,000 dilutions was used for immunoreaction with the anti-*Mmp*-aCPSF1 or anti-*Mmp*-aRpoD/L antiserum. Immune-active bands were visualized by an Amersham ECL Prime Western blot detection reagent (GE Healthcare). Protein quantitation was performed using Quantity One (Bio-Rad).

### Northern blot analysis and mRNA half-life determination

Northern blot was performed to determine various cellular RNA species and contents as described previously^55^. To measure the half-life of a specific transcript, a final concentration of 100 μg/ml actinomycin D (actD, MP Biomedicals) was added into the exponential cultures of *M. maripaludis* to stop transcription. At 0, 2, 4, 6, 8, and 10 min post-addition of actD, each 600 μl culture was collected and RNA was isolated. RNA content in each sample was quantified by Northern blot using the corresponding labeled DNA probe. A liner regression plot based on the average RNA contents from three batches of culture plotted against the sampling time was used for calculation the half-life of decay (t_1/2_) that was the time point when 50% RNA content retained.

To determine the bulk mRNA half-life, *M. maripaludis* was cultured to the exponential phase, and 30 μCi [5,6-^3^H]-Uridine (PerkinElmer) were added to isotope label the nascent transcripts. After a 10 min-labeling, actD was added and 600 μl culture was collected at 0, 2, 4, 6, 8, and 10 min post-addition of actD as described above. Then, 67 μl of 100% cold trichloroacetic acid (TCA) was added to each sample to precipitate the total RNA, and the mixture was incubated on ice for >30 min. A sample of 60 min post-addition of act D were used for counting the stable RNAs. RNA in each TCA mixture was precipitated by centrifuge at 12,000 g for 10 min and dissolved in 700 μl of Ecoscint A (National Diagnostics) for ^3^H isotope counting using liquid scintillator detector (PerkinElmer). The stable RNA ^3^H counts were first deducted from the total ^3^H ones at each sampling time point and the resultant count at 0 min post-addition of act D was recorded as 100% mRNA, and the residual mRNA content at each time point was plotted against the sampling time. Bulk mRNA half-life was calculated as described above.

### Electronic microscopy and motility assay

The mid-exponential cells of *M. maripaludis* strains S2 and ▽*aCPSF1* grown at 22°C were loaded onto carbon-Formvar-coated copper grids and washed with ddH_2_O and stained with uranyl acetate. Cells and archaella were observed under an LKB-V Ultratome (Jel1400) transmission electronic microscope. For mobility assay, cells were collected from 5 ml stationary phase culture grown at 22°C and resuspended gently in 200 μl of McF medium. 10 μl of cell suspension were inoculated in the semi-solid McF medium containing 0.25% (w/v) agar inside an anaerobic chamber. Plates were incubated at 22°C for 15 or 20 days in an anaerobic jar with Oxoid AnaeroGen (Thermo Scientific) to remove the oxygen.

### PALM imaging on the cellular location of *Mmp*-aCPSF1

Cell cultures of stains S2, *Mmp*-aCPSF1-mMaple and *Mmp*-aRpoL-mMaple (Supplementary Table 1) at earlier-(OD600≈0.2) or late-exponential phases (OD600≈0.7) grown at 37°C were collected by centrifugation, washed twice and suspended in an isotonic buffer (0.4 M NaCl, 60 mM NaHCO_3_ (pH 8.0)). Cells were then fixed by 0.1% poly-Lysine for 15 min at room temperature, washed three times with the isotonic buffer, and observed via PALM imaging. PALM imaging was obtained using a Nikon TiE inverted microscope equipped with a 100x oil-immersion objective (Nikon, PLAN APO, 1.49 NA) and an EMCCD camera (Andor-897) as described previously^56^. When PALM images obtained, the same fields were strained with 1% DAPI for 1 min and activated with a 405-nm laser to locate the nucleoid regions. The PALM images were constructed using Insight3 software, and data analysis was carried out using the custom-written Matlab scripts.

### Size Exclusion Chromatography

The exponential cells of *M. maripaludis* S2 culture were harvested and resuspended in the lysis buffer (50 mM Tris-HCl (pH 7.5), 150 mM NaCl, 0.05% NP-40 and 10% (v/v) glycerol), and then lysed by sonication and centrifuged at 12,000g for 30 min at 4°C. The supernatant was incubated at 37°C for 30 min with or without treatment of DNase I (100 U), RNase A (100 U) or Benzonase Nuclease (100 U). 500 μg of the pretreated supernatant were fractionated through the Superdex 200 10/300 GL column (GE Healthcare) and the molecular sizes of each fraction was estimated by gel filtration molecular weight markers (Kit No: MWGF1000 of Sigma-Aldrich), and each fraction was collected for protein identification by Western blot.

### Co-immunoprecipitation and Tandem Affinity Chromatography (TAP)

Association of *Mmp*-aCPSF1 with *Mmp*-aRNAP was detected by co-immunoprecipitation and Tandem Affinity Chromatography. To perform anti-Flag co-immunoprecipitation, prewashed Anti-FLAG M2 Magnetic Beads (Sigma) was added to 5 mg pretreated cell extract and incubated 8 h at 4°C with gently shaking. The antigen bound Magnetic Beads were washed five times with the lysis buffer and eluted by 2.5 volume of 3×FLAG peptide (150 ng/µl, Sigma) for 3h at 4°C with gently shaking. For His-tag affinity chromatography, ∼20 mg cell extract was purified through a His-Trap HP column as described above. For TAP, ∼20 mg cell extract was first purified by His-tagged affinity chromatography, and then the product was concentrated and removed of imidazole using Amicon Ultrafra-30 concentrators (Millipore). The concentrated product was then immunoprecipitated by the same procedure as for anti-Flag co-immunoprecipitation. All of the CoIP, His-tagged affinity chromatography and TAP products were separated on a 12% SDS-PAGE gel and identified by Western blot.

### Chromatin Immunoprecipitation (ChIP) coupled quantitative PCR (ChIP-qPCR)

To obtain the chromosomal occupancy of *Mmp*-aCPSF1 and *Mmp-aRpoL*, ChIP assays were performed for strains *Mmp*-aCPSF1-F and *Mmp*-aRpoL-HF as described previously^57–59^ with some modifications. Cells of *M. maripaludis* S2 were used as mock controls. Middle exponential cells of ∼0.5-1×10^9^ cells ml^-1^ were formaldehyde fixed and lysed. The chromosomal DNA in the lysed supernatant was sheared by sonication to an average size of 200-500 bp using Bioruptor UCD300 (Diagenode) before incubation overnight at 4°C with Anti-FLAG M2 Magnetic Beads (Sigma). The beads were then washed, and the DNA-protein complexes were eluted by the ChIP elution buffer (10 mM Tris-HCl (pH 8.0), 1 mM EDTA, 1% SDS) at 65°C for 30 min with shaking. Reverse crosslinking was then performed by treated with 0.05 mg/ml RNase A and 0.5 mg/ml proteinase K at 37°C for 2–4 h followed by overnight incubation at 65°C. DNA fragments were purified using MiniElute columns (Qiagen) and quantified using the Qubit dsDNA HS kit (Life Technologies). PCR and qPCR were performed to determine the DNA species and the enrichment by *Mmp*-aCPSF1 or *Mmp*-aRpoL as described previously^58, 60^. For PCR amplification, 1 μL each of input and IP or mock-IP DNA sample was used in a 25 μl reaction mix containing 400 nM of the corresponding oligonucleotide primer (Supplementary Table 2). For qPCR analysis, similar reaction mixture of PCR except for 1×SYBR Green Real-time PCR Master Mix (Toyobo, Tokyo, Japan) was used and performed at Mastercycler EP realplex2 (Eppendorf, Germany). To estimate the copy numbers of aCPSF1 or RNAP captured genes, standard curves of the corresponding genes were generated using 10-fold serially diluted PCR product as templates. The enrichment percentage of each gene was calculated as the copies in the sample of ChIP divided by that in the input and then times 100. The enrichment fold was calculated as the ChIP enriched percentage over that of the mock.

## Acknowledgments

The authors thank Prof. William B. Whitman at the University of Georgia providing strain *Methanococcus maripaludis* S2 and plasmids pIJA03 and pMEV2, Prof Tao Yong at Institute of Microbiology, CAS providing *Escherichia coli* BW25113 and plasmid pSB1s, and Prof. Fan Bai at Peking University to assist PALM imaging. This work is supported by the National Natural Science Foundation of China under grant nos. 31430001, 91751203, and 31670049. The authors also thank LetPub (www.letpub.com) for linguistic assistance on the manuscript preparation.

## Author contributions

L.Y and J.L designed and performed the genetic and biochemical experiments. J.L and X.Z.D conceptualized the experiments and acquired funding. L.Y and L.Q performed the transcriptomics assays. B.Z and F.Q.Z performed the bioinformatics analyses. L.L cultured strains. J.L and X.D wrote the manuscript and all of the authors approved the final manuscript.

**Authors declare that they do not have any competing interests**

**All data are available in main text or supplementary materials.**

**List of Supplementary Materials**:

Supplementary Fig. 1-9;

Supplementary Table 1-2;

Supplementary Datasets 1-4.

