## Supplementary Information for "aCPSF1 controlled archaeal transcription termination: a prototypical eukaryotic model"

**This file contains:**

Supplementary Figs. 1 to 9

Supplementary Tables 1 to 2

Caption for Supplementary Dataset 1: Differential transcription of *M. maripaludis* S2 vs *Mmp-aCPSF1* depletion mutant (▽*aCPSF1*).

Caption for Supplementary Dataset 2: Defined transcriptional units (TUs) in the *M. maripaludis* S2 transcriptome and calculated transcriptional read-through (TRT).

Caption for Supplementary Dataset 3: Term-seq identified primary transcriptional end sites (TESs) in *M. maripaludis* S2.

Caption for Supplementary Dataset 4: Prevalence of the uridine-rich sequences in the IGRs among Archaea.

### Supplementary Figures

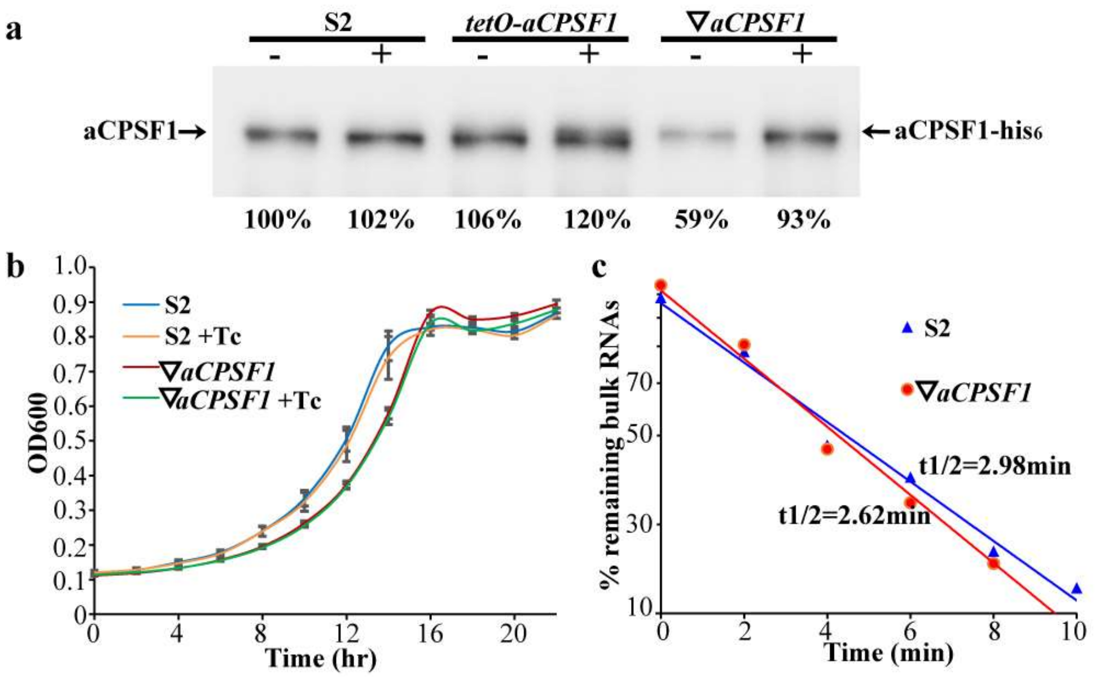


**Supplementary Figure 1. Depleted expression of *Mmp-aCPSF1* showed slight effects the growth and cellular RNA lifespan of *M. maripaludis* S2 at 37°C. a,** Western blot detected the protein abundance (percentages by referenced to that of lane 1 shown beneath) of *Mmp*-aCPSF1 in the presence (+) or absence (–) of 100 µg/ml tetracycline (Tc) in S2 (wild-type), *tetO-aCPSF1*, and ∇*aCPSF1* at 37°C. aCPSF1 and aCPSF1-His$_6$ flanked the gel pointed the indigenous and *hpt*-site inserted *Mmp*-aCPSF1, respectively. **b,c,** *Mmp-aCPSF1* depletion did not affect growth (**b**) and the life-span of total mRNA in 37°C cultured cells (**c**). Half-lives of total mRNA are calculated from the regression curve of residual mRNA by quantifying the [$^3$H]-uridine signal attenuation as described in Methods. Experiments were performed on three batches of culture, and standard deviations were shown.

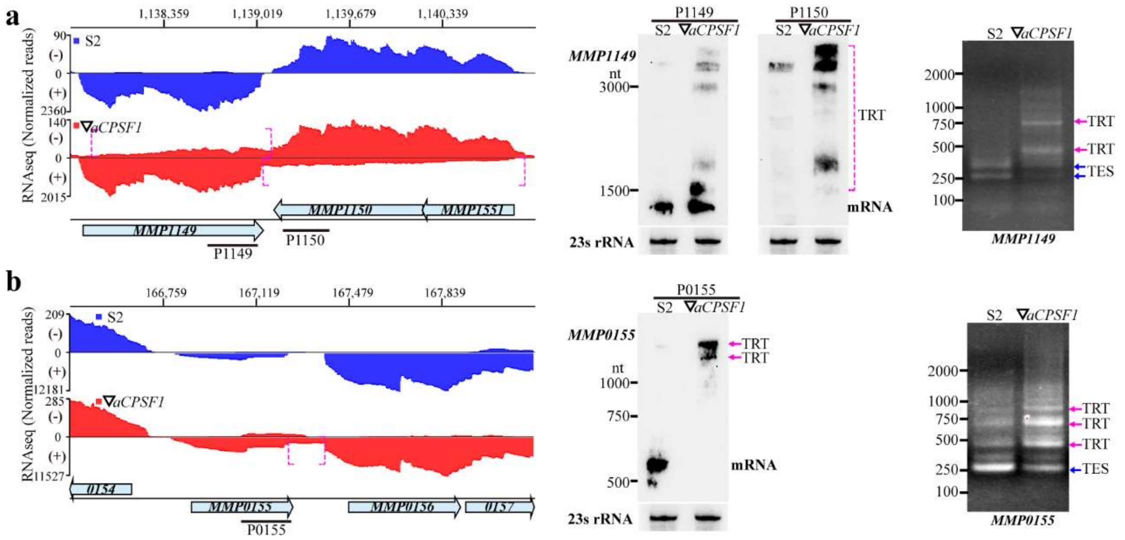


**Supplementary Figure 2. Northern blot and 3′RACE verifications of RNA-seq detected TRT in the *Mmp-aCPSF1* depletion mutant.** Similar to Fig. 2a, *MMP1149*, encoding 3-isopropylmalate dehydratase, and the antisense *MMP1150* for uroporphyrinogen decarboxylase (**a**), and *MMP0155* encoding a putative RNA-binding protein (**b**) that exhibited 3′-end extension (bracketed with magenta line) upon *Mmp-aCPSF1* depletion were selected for verification. Left panels show the RNA-seq reads mapped to the corresponding genes in strains S2 (blue) and ∇*aCPSF1* (red). Numbers on the top indicate the nucleotide sites of mapped genomic regions, and bullets represent gene orientations. Middle panels show the transcript with natural termination sites (TES) and the TRT (magenta arrows or frame) transcripts detected by Northern blot. Right panels show 3′RACE amplified products with natural TES (blue arrows) and TRT (magenta arrows) in S2 and ∇*aCPSF1*, respectively. 3′RACE assayed 3′-end nucleotide sequences are shown in Supplementary Fig. 5.

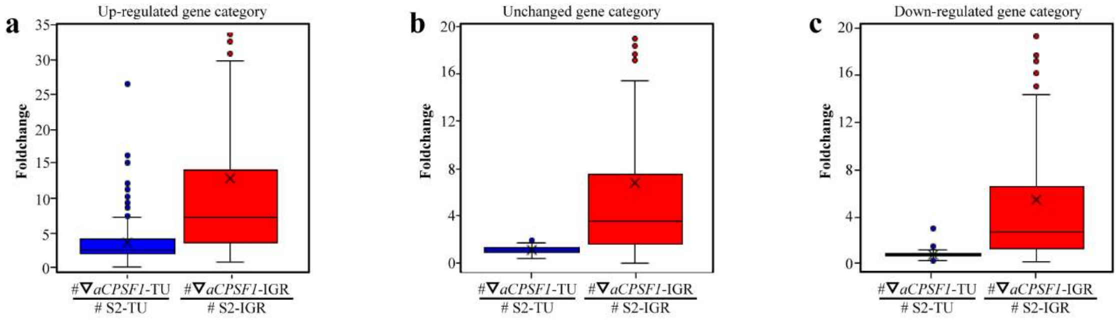


**Supplementary Figure 3. *Mmp-aCPSF1* depletion caused higher transcription increasing in IGR than the TU body**. Based on the transcription changes upon *Mmp-aCPSF1* depletion, TUs are first classified into three categories as Up-regulated (**a**), unchanged (**b**), and down-regulated (**c**). Boxplots show the statistics of the fold changes in IGRs and the associated TU bodies, which displays the fold change values among 50% genes (inside the box) and the medians (line inside the box).

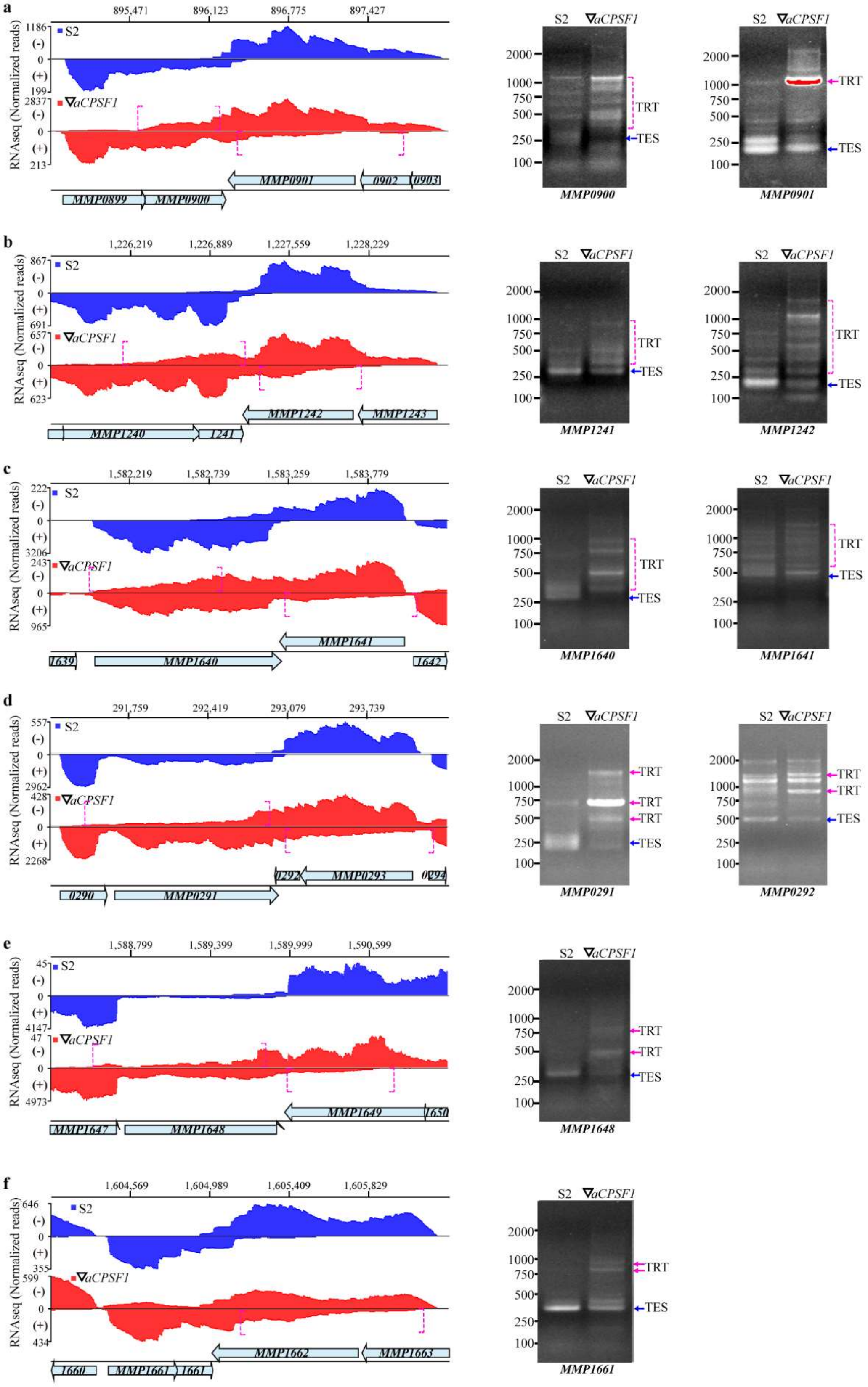
a
895,471
896,123
896,775
897,427
RNAseq (Normalized reads)
S2
∇aCPSF1
MMP0899
MMP0900
MMP0901
0902
0903
MMP0900
MMP0901
TRT
TES
b
1,226,219
1,226,889
1,227,559
1,228,229
MMP1240
1241
MMP1242
MMP1243
MMP1241
c
1,582,219
1,582,739
1,583,259
1,583,779
1639
MMP1640
MMP1641
1642
d
291,759
292,419
293,079
293,739
0290
MMP0291
0292
MMP0293
0294
MMP0292
e
1,588,799
1,589,399
1,589,999
1,590,599
MMP1647
MMP1648
MMP1649
1650
f
1,604,569
1,604,989
1,605,409
1,605,829
1660
MMP1661
1661
MMP1662
MMP1663
2000
1000
750
500
250
100

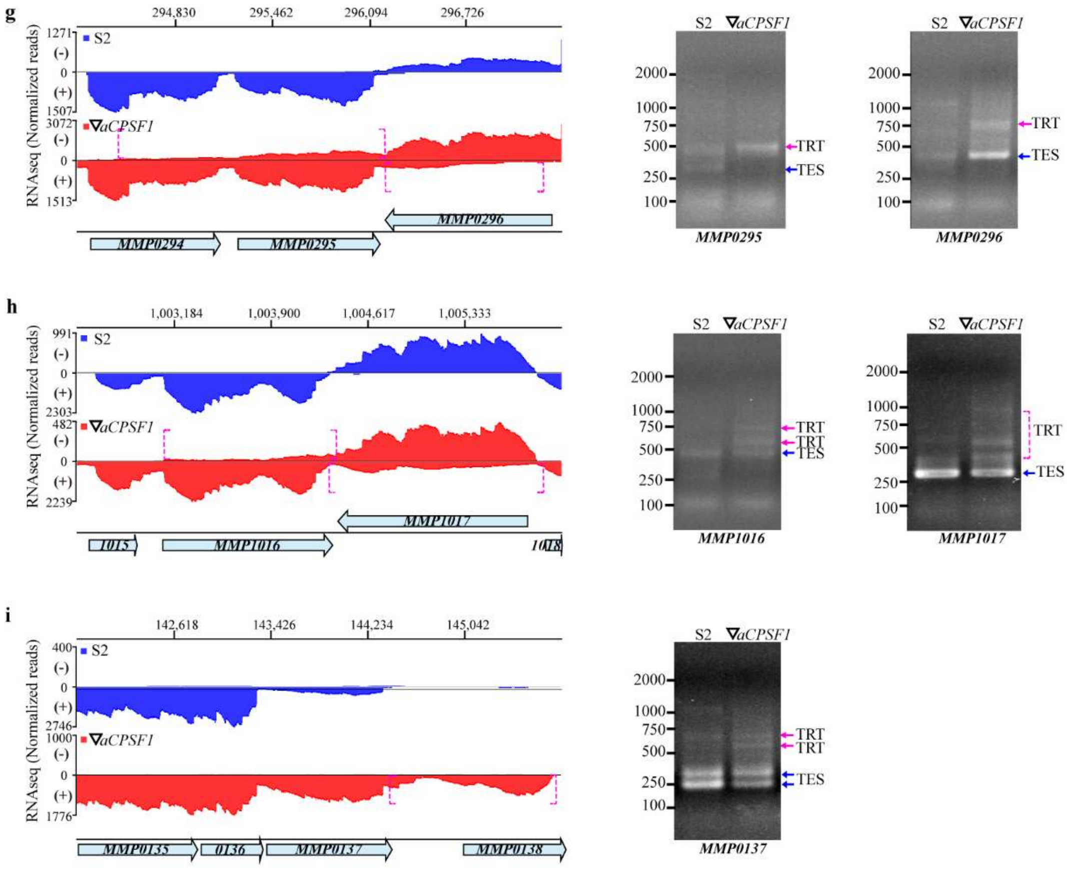


**Supplementary Figure 4. 3′RACE validation of *Mmp-aCPSF1* depletion caused TRTs in fifteen genes.** Using the same approach as in Fig. 2a and supplementary Fig. 2, additional fifteen representative transcripts that exhibited 3′-end extension (dotted magenta brackets) in ∇*aCPSF1* were selected for experimental validation. *MMP0900* and *MMP0901* encode tRNA 2-selenouridine synthase and a small GTP-binding protein, respectively (**a**); *MMP1241* and *MMP1242* encode a hypothetical protein and quinolinate synthetase subunit A, respectively (**b**); *MMP1640* and *MMP1641* encode S-adenosylmethionine synthetase and soluble P-type ATPase (**c**); *MMP0291, MMP0292* (**d**)*, MMP1648, MMP1449* (**e**), *MMP1661* and *MMP1663* (**f**) all encode hypothetical proteins; *MMP0295* and *MMP0296* encode homoserine kinase and nonsense-mediated mRNA decay protein (**g**); *MMP1016* and MMP1017 encode a hypothetical protein and aspartate kinase, respectively (**h**); and *MMP0137* encodes deoxyhypusine synthase (**i**). Left panels show the RNA-seq reads mapped to the corresponding genes in strains S2 (blue) and ▽*aCPSF1* (red), respectively. Numbers on the top indicate the nucleotide sites of mapped genomic regions, and bullets represent gene orientations. Right panels show 3′RACE amplified products with natural termination sites (TES, blue arrows) and TRT (magenta arrows) of the indicated genes beneath the gel. 3′RACE assayed nucleotide sequences of the transcript 3′-ends are shown in Supplementary Fig. 5.

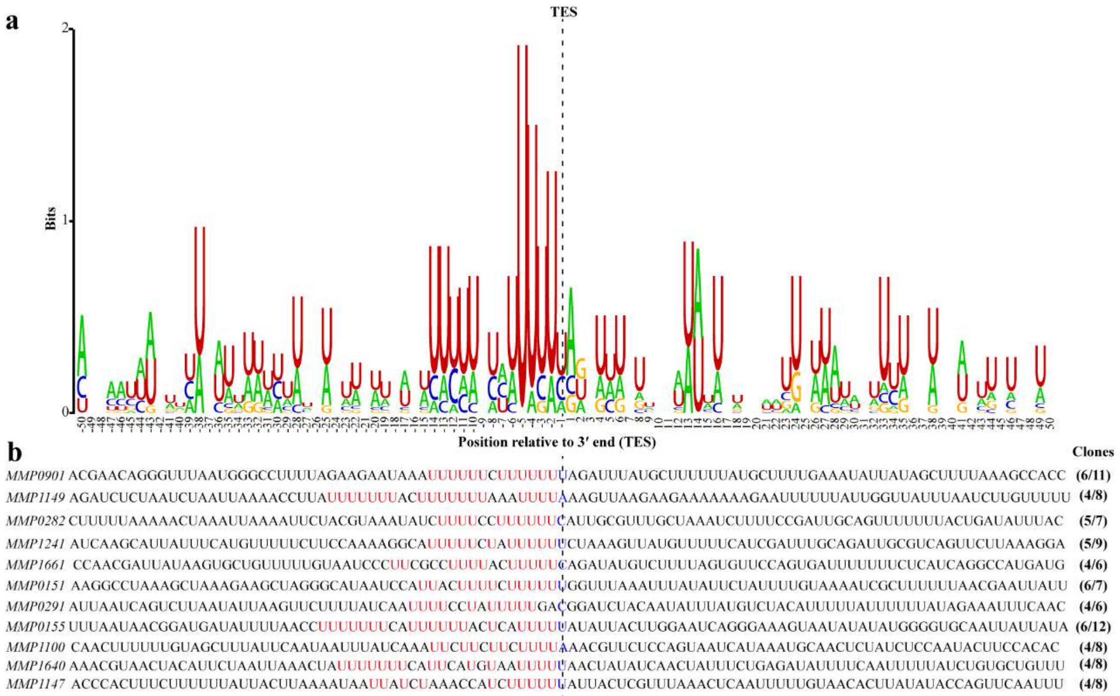


**Supplementary Figure 5. 3′RACE determined terminator motifs**. Using the MEME algorithm, a terminator motif of uridine-rich tract (**a**) was searched upstream 3′RACE captured 3′-ends of transcripts (TES, dot line) among 11 transcription units (**b**). The sequence logos were generated by WebLogo. Similar terminator motif was also found in the Term-seq data (Fig. 4a). The 50 nt sequence each upstream and downstream the TES determined in the 11 TUs are shown in b, in which uridine-rich stretches are in red. Inside parenthesis indicate the sequenced clone numbers of 3′RACE amplified fragments, and the former indicate clone numbers that matching the same TES sites detected by Term-seq.

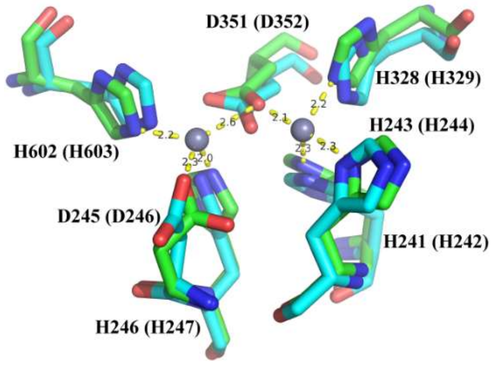


**Supplementary Figure 6. Predicted catalytic center of *Mmp*-aCPSF1**. Using mm_0695 (PDB ID: 2XR1), an aCPSF1 from *Methanosarcina mazei* as template, the protein structure of *Mmp*-aCPSF1 (MMP0694) was homology-modelled using SWISS-MODEL (https://swissmodel.expasy.org/). The catalytic center with two zinc ions (light blue spheres) and seven conserved residues (sticks of green for mm_0695 and cyan for MMP0694) is shown. Inside the parentheses indicate the residue numbers of *Mmp*-aCPSF1. Dot yellow lines show the predicted hydrogen-bond interactions between zinc ions and residues.

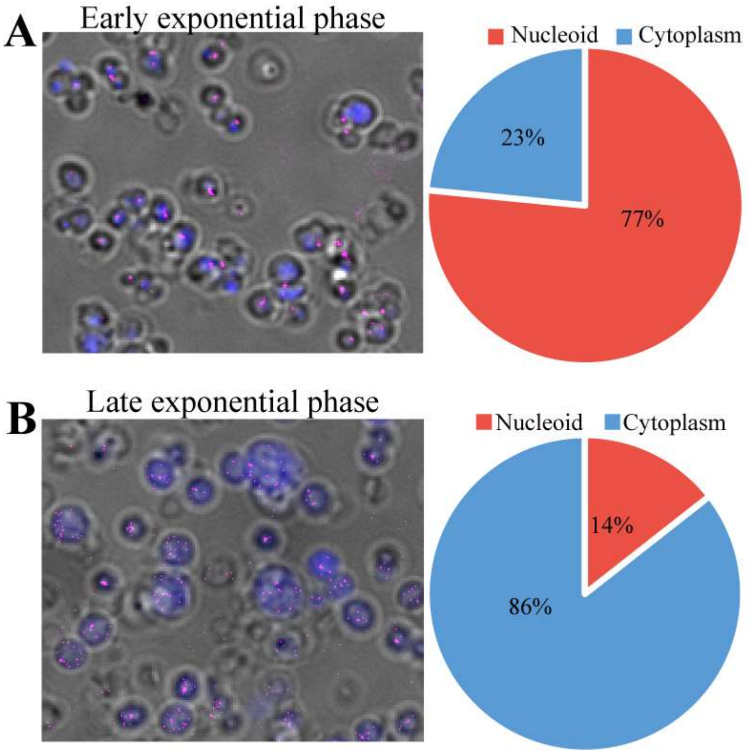


**Supplementary Figure 7. Variable cellular location of *Mmp*-aCPSF1 protein during growth.** Using the approach described in Fig. 5d and in Methods, the representative super-resolution PLAM microscopic images show the cellular locations of *Mmp*-aCPSF1 protein in the early (**a**) and later exponential (**b**) growth phases were captured. By observing at least six microscopic fields, cells with the mMaple3-fused aCPSF1 protein in the nucleoid and cytoplasm were counted and the average percentages were calculated (right panel).

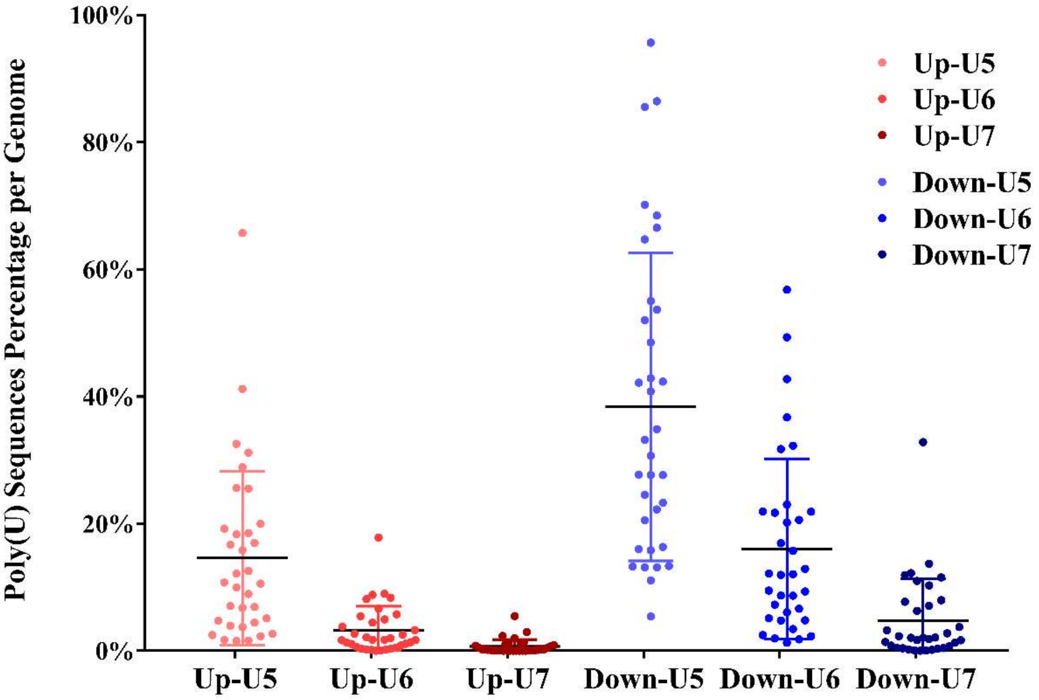


**Supplementary Figure 8. Enrichment of the uridine-rich sequences in the IGRs of archaeal genome.** The uridine-rich tract (U5, U6 and U7) sequences were searched at the 200 nt upstream (up) and downstream (Down) the predicted stop codons of annotated ORFs in the representative species from all four archaeal superphyla shown in Supplementary Dataset 4. The Poly(U) sequence percentage was calculated by the uridine-rich tract numbers over the gene numbers in each selected genome. Black lines refer to the percentage medians and the colored lines indicate the lower to the upper quartile, respectively. U5, U6 and U7 indicate 5, 6 and 7 successive uredines, respectively.

```
Lokiarch_44440  M  S  F  E  L  E  L  K  R  I  R  D  E  I  V  Q  N  L  P  P  E  I  T  V  T  K  I  E  F  E
Original 1      ATGAGTTTTGAATTAGAACTTAAAAGAATTAGAGACGAAATTGTCCAGAATCTACCCCCCGAAATTACTGTTACAAAAATCGAATTTGAG
Optimized       ATGTCATTTGAATTAGAACTTAAAAGAATTAGAGATGAAATAGTTCAAAATTTGCCACCAGAAATAACAGTTACTAAAATAGAATTTGAA
Lokiarch_44440  G  P  E  I  A  V  Y  S  E  S  E  D  V  A  A  I  E  T  S  S  T  L  K  D  L  A  K  V  M  R
Original_91     GGTCCAGAAATTGCTGTTTATTCAGAGAGTGAGGATGTTGCCGCTATTGAAACCTCTTCTACCTTAAAAGATTTAGCGAAGGTGATGAGA
Optimized       GGACCAGAAATTGCAGTTTATTCAGAAAGTGAAGATGTTGCAGCAATTGAAACTTCATCTACTTTAAAGGATCTTGCAAAAGTTATGAGA
Lokiarch_44440  K  R  V  V  F  R  W  N  V  D  K  R  K  D  P  A  E  T  K  D  Y  I  M  N  L  I  S  K  D  A
Original_181    AAAAGAGTTGTGTTTAGGTGGAACGTAGATAAACGCAAAGACCCAGCAGAAACTAAAGACTATATAATGAACCTAATCAGCAAAGATGCT
Optimized       AAAAGAGTAGTATTCAGATGGAATGTTGATAAAAGGAAAGATCCTGCTGAAACTAAAGATTACATTATGAATCTTATTTCAAAAGATGCA
Lokiarch_44440  E  I  G  E  I  T  F  D  H  T  R  G  E  V  I  I  E  S  G  K  P  G  L  V  I  G  K  K  G  I
Original_271    GAAATTGGCGAAATAACCTTCGATCACACCCGTGGCGAAGTTATCATCGAATCAGGAAAACCAGGACTAGTAATAGGTAAAAAAGGAATT
Optimized       GAAATAGGTGAAATAACTTTTGATCACACTAGAGGTGAAGTAATTATTGAATCAGGAAAACCAGGATTAGTAATTGGTAAAAAAGGTATA
Lokiarch_44440  N  L  K  E  I  R  T  N  T  F  W  Q  P  K  T  I  R  T  P  P  L  P  S  R  T  I  Q  L  I  R
Original_361    AATTTAAAAGAAATTAGAACAAACACATTCTGGCAGCCAAAAACTATCAGAACACCACCTCTACCTTCTAGAACAATTCAATTAATTAGA
Optimized       AATCTTAAAGAAATTAGAACAAATACATTTTGGCAGCCTAAAACAATAAGAACCCCACCTTTACCTTCTAGGACTATTCAGTTAATAAGA
Lokiarch_44440  G  M  L  K  K  E  R  Q  T  Q  K  D  I  L  L  E  I  G  K  R  I  H  R  P  A  L  F  N  N  L
Original_451    GGTATGCTTAAAAAAGAAAGACAAACACAGAAAGACATTCTTTTAGAAATAGGAAAAAGAATTCACAGACCTGCACTATTTAACAATTTA
Optimized       GGAATGCTTAAAAAAGAAAGACAAACACAGAAAGATATTCTTTTAGAAATAGGTAAAAGAATTCATAGACCAGCATTATTCAATAACTTA
Lokiarch_44440  N  I  R  M  N  A  L  G  G  F  R  E  V  G  R  S  C  I  L  M  Q  T  R  D  S  N  V  L  L  D
Original_541    AATATAAGAATGAATGCATTAGGGGGTTTTCGCGAAGTTGGGCGATCCTGTATATTAATGCAGACGAGAGATAGTAATGTTTTACTAGAC
Optimized       AATATTAGAATGAATGCTTTAGGAGGTTTTAGAGAAGTAGGTAGATCATGCATTTTAATGCAGACTAGAGATTCAAATGTTTTATTAGAT
Lokiarch_44440  V  G  L  N  V  G  N  P  N  D  R  F  P  Y  F  F  V  P  Q  F  S  I  R  D  L  D  A  V  I  I
Original_631    GTGGGATTGAATGTAGGGAATCCGAATGACAGGTTTCCTTATTTCTTTGTCCCACAATTTTCCATTCGTGATCTTGATGCTGTAATCATT
Optimized       GTAGGTCTTAATGTTGGAAATCCAAATGATAGATTTCCATACTTTTTCGTTCCTCAATTTTCTATTAGGGATCTTGATGCAGTAATAATT
Lokiarch_44440  S  H  S  H  L  D  H  C  G  V  V  P  F  L  Y  K  Y  G  Y  R  G  P  I  Y  C  T  L  P  T  R
Original_721    TCACACTCGCACCTTGATCATTGTGGTGTTGTTCCCTTTCTATATAAATATGGTTATCGTGGTCCAATTTATTGTACATTACCTACAAGA
Optimized       AGTCATTCTCACCTTGATCACTGCGGAGTTGTTCCATTTTTATATAAATATGGTTATAGAGGACCAATCTATTGTACACTTCCAACAAGA
Lokiarch_44440  N  L  S  T  M  L  Q  L  D  F  I  Q  I  C  D  K  E  G  T  P  S  P  Y  S  K  R  D  V  K  N
Original_811    AATTTATCAACAATGTTACAGCTTGATTTCATTCAAATTTGTGACAAAGAAGGAACTCCATCACCATATTCAAAAAGAGATGTTAAAAAT
Optimized       AACCTTTCAACAATGTTACAGTTAGATTTTATTCAAATATGTGATAAAGAAGGAACTCCAAGTCCATATAGTAAAAGAGATGTTAAAAAT
Lokiarch_44440  A  V  L  H  T  I  P  L  S  W  G  K  V  T  D  I  A  P  D  I  K  L  T  L  H  N  S  G  H  I
Original_901    GCAGTACTTCATACAATTCCCCTCTCTTGGGGTAAAGTTACTGACATTGCTCCGGATATAAAACTTACACTTCATAATTCGGGTCATATT
Optimized       GCAGTATTACATACCATACCATTAAGTTGGGGTAAAGTTACAGATATTGCTCCTGATATAAAATTAACATTACATAATAGTGGACACATT
Lokiarch_44440  L  G  S  S  M  I  H  L  H  F  G  K  G  D  Y  N  F  V  Y  T  G  D  F  K  Y  Q  K  T  R  L
Original_991    TTAGGATCTTCTATGATACATTTACATTTTGGGAAGGGAGATTATAATTTTGTGTATACAGGGGATTTTAAATATCAAAAAACTCGATTG
Optimized       TTGGGTTCATCAATGATTCACTTACACTTTGGTAAAGGAGATTATAATTTTGTTTACACAGGAGATTTTAAATACCAAAAAACAAGGTTA
Lokiarch_44440  L  E  K  A  T  V  K  F  P  R  V  E  G  L  L  I  E  A  T  Y  G  G  P  Q  D  R  I  P  S  R
 Original_1081  CTAGAAAAAGCCACAGTTAAATTTCCAAGAGTTGAAGGCCTTTTAATTGAAGCTACTTATGGTGGGCCTCAAGATCGCATCCCAAGCAGA
Optimized       TTAGAAAAAGCAACTGTAAAATTTCCAAGAGTTGAAGGACTTTTAATAGAAGCAACATATGGTGGTCCTCAAGATAGAATTCCTTCTAGA
Lokiarch_44440  Q  E  S  E  K  E  L  K  Q  I  L  N  S  T  L  K  R  G  G  K  V  L  I  P  V  L  A  V  G  R
 Original_1171  CAAGAATCTGAAAAGGAGTTAAAACAGATTTTAAATTCCACACTCAAAAGAGGTGGAAAAGTTCTAATACCTGTTCTGGCTGTAGGACGT
Optimized       CAAGAATCTGAAAAAGAACTCAAACAAATTCTTAATTCAACTCTTAAAAGAGGTGGAAAAGTTTTAATTCCAGTATTAGCAGTAGGAAGA
Lokiarch_44440  A  Q  E  L  L  I  V  L  E  E  Y  I  S  S  G  F  I  D  K  V  P  V  Y  I  D  G  L  I  S  E
 Original_1261  GCTCAGGAATTATTAATTGTTTTAGAAGAATACATCTCATCAGGATTTATTGATAAAGTACCAGTCTATATTGATGGTCTAATAAGTGAA
Optimized       GCTCAGGAATTATTAATTGTTTTAGAAGAATATATTTCATCAGGTTTTATTGATAAAGTTCCAGTTTATATTGATGGATTAATTTCAGAA
Lokiarch_44440  A  T  A  I  H  T  A  N  P  D  F  L  S  S  D  L  R  E  K  I  L  H  Q  G  K  N  P  F  L  S
 Original_1351  GCCACCGCAATCCACACGGCCAATCCTGATTTTCTGAGCAGTGATTTAAGAGAAAAAATACTTCATCAAGGGAAAAATCCATTTTTAAGC
Optimized       GCAACAGCAATTCATACGGCTAATCCTGACTTTTTAAGTTCTGACTTAAGAGAAAAAATATTACACCAAGGAAAAAATCCATTTTTAAGT
Lokiarch_44440  D  F  F  E  T  V  S  S  Q  D  E  R  A  D  I  I  G  G  G  P  C  I  I  L  A  T  S  G  M  L
 Original_1441  GACTTTTTTGAGACCGTTTCATCTCAAGATGAGAGAGCTGACATAATTGGGGGAGGTCCATGTATAATACTAGCTACTAGTGGCATGCTA
Optimized       GATTTCTTTGAAACTGTTTCTTCTCAGGATGAAAGAGCAGATATTATAGGAGGAGGACCATGTATAATTTTAGCAACATCAGGAATGTTG
Lokiarch_44440  I  G  G  P  S  V  H  Y  L  K  A  L  A  E  D  S  K  N  T  L  I  F  V  S  Y  Q  V  S  G  T
 Original_1531  ATTGGAGGACCTTCAGTTCATTACTTAAAAGCTCTAGCAGAGGATTCAAAAAACACATTGATTTTCGTATCTTATCAAGTTTCAGGGACT
Optimized       ATAGGAGGACCTAGTGTTCATTACTTAAAAGCTTTAGCAGAAGATTCAAAAAATACATTAATTTTTGTAAGTTACCAAGTTTCAGGAACA
Lokiarch_44440  L  G  S  R  I  I  R  G  F  R  E  F  N  Y  I  D  W  K  G  R  T  Q  L  V  K  I  G  L  K  V
 Original_1621  TTAGGAAGCAGGATAATTCGTGGGTTTAGAGAATTCAATTATATAGATTGGAAAGGTCGAACACAGCTAGTAAAAATAGGTTTAAAAGTA
Optimized       TTAGGATCTAGAATAATAAGAGGTTTTAGAGAATTCAATTACATTGATTGGAAAGGAAGAACACAGTTGGTTAAAATAGGACTTAAAGTA
Lokiarch_44440  F  T  L  E  G  F  S  G  H  S  S  R  S  Q  I  S  Q  F  L  R  R  I  Q  P  R  P  K  V  V  I
 Original_1711  TTTACTCTTGAAGGTTTTAGTGGACATTCATCTCGTAGCCAAATTTCTCAATTTCTGAGAAGAATTCAGCCAAGACCAAAGGTAGTTATA
Optimized       TTCACACTTGAAGGATTTTCAGGACACTCATCAAGATCACAGATTTCACAGTTTTTAAGAAGAATTCAGCCTAGACCTAAAGTTGTAATT
Lokiarch_44440  V  N  H  G  E  E  S  K  C  V  S  L  S  T  M  I  H  K  K  L  R  K  S  T  K  S  P  K  N  L
 Original_1801  GTAAATCATGGAGAAGAAAGTAAATGCGTAAGTTTATCAACAATGATACATAAAAAGCTAAGGAAATCAACTAAAAGTCCTAAAAATCTT
Optimized       GTTAATCACGGAGAAGAATCTAAATGTGTATCTTTAAGTACAATGATTCATAAAAAATTAAGAAAAAGTACTAAATCACCTAAAAACTTA
Lokiarch_44440  E  V  V  L  L  K  *
Original_  1891 GAAGTTGTATTACTTAAATAA
Optimized       GAAGTAGTTCTTTTAAAATAA
```

```
CENSYa_1545   M  Q  R  R  Q  Q  Q  K  E  A  P  S  N  Q  N  I  M  A  T  I  L  G  S  I  P  R  E  A  G  V
Original 1    ATGCAACGCAGACAGCAGCAAAAGGAGGCGCCGAGCAACCAGAACATAATGGCGACCATCCTTGGCAGCATCCCGAGGGAGGCCGGTGTG
Optimized     ATGCAGAGAAGACAACAGCAGAAAGAAGCACCAAGTAATCAAAATATTATGGCAACAATTTTAGGATCAATACCTAGAGAAGCAGGAGTT

CENSYa_1545   T  K  I  E  Y  E  G  P  R  I  A  L  Y  T  N  T  P  R  Y  L  L  E  H  N  E  I  I  S  N  L
Original 91   ACAAAGATAGAATACGAGGGCCCGCGCATAGCGCTGTATACGAATACGCCGCGATACCTCCTGGAGCACAACGAGATAATATCGAACCTG
Optimized     ACTAAAATTGAATATGAAGGACCAAGAATTGCTCTTTATACAAATACACCAAGATATCTTTTAGAACACAATGAAATAATTTCAAATCTT

CENSYa_1545   V  N  V  I  K  K  R  I  I  V  R  I  D  E  S  V  R  K  K  E  E  D  A  R  K  M  L  S  K  L
Original 181  GTAAACGTGATAAAAAAAAGAATCATAGTAAGGATAGACGAATCGGTCCGGAAAAAGGAGGAGGACGCGCGGAAGATGCTCTCAAAGCTC
Optimized     GTAAATGTTATCAAAAAACGAATTATTGTAAGAATTGATGAATCTGTTAGAAAAAAGGAAGAAGATGCAAGAAAAATGTTAAGTAAATTA

CENSYa_1545   V  P  A  E  A  K  L  Q  G  T  F  F  D  T  T  T  G  E  V  S  L  E  A  K  R  P  W  L  L  Q
Original 271  GTCCCTGCCGAGGCGAAGCTGCAGGGCACATTCTTTGACACCACGACGGGCGAGGTCTCCCTGGAGGCCAAGAGGCCGTGGCTGCTCCAG
Optimized     GTACCAGCTGAAGCTAAACTTCAAGGAACTTTCTTTGATACAACTACAGGAGAAGTAAGTTTAGAAGCAAAAAGACCATGGTTACTTCAA

CENSYa_1545   R  N  S  A  E  F  S  H  A  D  V  S  E  K  I  G  W  T  L  R  I  R  K  A  T  T  S  Q  S  S
Original 361  CGCAACTCAGCCGAGTTCAGCCACGCCGATGTGAGCGAAAAGATAGGCTGGACGCTCCGGATTAGAAAGGCCACCACCAGCCAGTCGAGC
Optimized     AGAAATTCAGCTGAATTTTCACACGCTGATGTTTCTGAAAAAATTGGATGGACATTAAGGATTAGAAAAGCAACAACAAGTCAGTCTTCT

CENSYa_1545   T  M  Q  V  I  N  R  T  L  R  A  S  S  I  E  R  G  K  Q  L  K  Q  I  G  D  D  I  F  R  P
Original 451  ACCATGCAGGTGATAAACCGCACATTGCGCGCATCGTCTATCGAGCGCGGCAAGCAGCTCAAGCAGATAGGCGACGACATATTCCGGCCC
Optimized     ACTATGCAAGTTATTAATAGAACCTTGAGGGCATCATCAATAGAAAGAGGTAAACAATTGAAACAAATAGGAGATGATATTTTTAGACCA

CENSYa_1545   K  L  A  T  R  S  E  I  S  L  T  A  L  G  G  F  G  Q  V  G  R  S  C  M  L  L  S  T  L  D
Original 541  AAGCTGGCCACCCGCTCGGAGATCTCGCTGACAGCGCTTGGCGGCTTCGGCCAGGTGGGCCGCTCGTGCATGCTGCTCTCTACTCTTGAC
Optimized     AAATTAGCAACTAGATCAGAAATTTCTTTAACAGCATTAGGTGGATTTGGACAAGTAGGTAGATCATGTATGCTTTTATCAACTCTTGAT

CENSYa_1545   S  K  V  L  V  D  C  G  V  N  P  G  A  A  H  P  S  E  S  Y  P  R  L  D  W  A  G  I  T  L
Original 631  AGCAAGGTTCTAGTCGACTGCGGGGTAAACCCGGGGGCGGCGCACCCGTCGGAGTCGTATCCGCGGCTCGACTGGGCGGGCATAACACTC
Optimized     AGTAAAGTTTTAGTAGATTGCGGAGTTAATCCAGGAGCAGCACACCCTAGTGAAAGTTATCCTAGACTTGATTGGGCAGGAATAACACTT

CENSYa_1545   D  D  L  D  A  V  V  I  G  H  A  H  L  D  H  T  G  F  L  P  V  L  A  K  Y  G  Y  R  G  P
Original 721  GACGACCTTGACGCGGTGGTCATAGGGCATGCGCATTTAGACCACACGGGCTTTCTGCCCGTTCTAGCAAAGTACGGGTACAGGGGCCCG
Optimized     GATGACTTAGATGCAGTTGTTATTGGACACGCACACTTAGATCATACTGGATTTTTACCAGTTTTAGCAAAATACGGATATAGAGGACCA

CENSYa_1545   I  Y  C  T  E  P  T  L  P  M  M  N  L  I  Q  L  D  A  I  K  V  A  T  A  Q  G  R  V  P  V
Original 811  ATATACTGTACAGAGCCGACGCTCCCCATGATGAACCTGATACAGCTCGATGCAATCAAGGTGGCTACAGCCCAGGGCAGGGTGCCCGTG
Optimized     ATATATTGTACAGAACCAACGTTACCTATGATGAATTTGATTCAATTAGATGCTATAAAAGTAGCAACAGCACAGGGTAGAGTTCCAGTT

CENSYa_1545   Y  A  E  R  D  V  R  Q  I  M  R  Q  A  I  T  L  P  Y  G  T  V  T  D  I  S  P  D  I  K  L
Original 901  TACGCCGAGCGCGACGTCCGGCAGATAATGAGGCAGGCGATCACTCTGCCCTATGGGACTGTCACCGATATCTCCCCGGATATCAAACTG
Optimized     TACGCAGAAAGAGATGTTAGACAGATTATGAGACAAGCAATTACTCTTCCATATGGAACTGTAACTGATATATCACCAGATATAAAATTA

CENSYa_1545   V  L  A  N  A  G  H  I  L  G  S  A  L  C  H  F  H  I  G  S  G  D  H  N  F  V  Y  S  G  D
Original 991  GTGCTGGCAAACGCGGGGCACATACTGGGATCCGCGCTCTGCCACTTCCACATAGGCAGCGGCGACCACAACTTTGTCTATTCAGGGGAC
Optimized     GTTTTAGCAAATGCAGGTCACATTTTAGGTTCTGCTCTTTGTCATTTTCACATAGGTTCAGGAGACCATAACTTTGTATACTCAGGAGAT

CENSYa_1545    I  K  F  G  K  S  I  L  F  E  A  A  S  W  N  F  P  R  V  E  T  L  L  I  E  S  T  Y  G  A
Original 1081 ATAAAGTTCGGCAAGAGCATACTATTCGAGGCGGCAAGCTGGAACTTTCCGCGCGTGGAGACTCTATTGATAGAGAGCACGTACGGGGCT
Optimized     ATAAAATTTGGAAAATCTATATTATTTGAAGCAGCAAGTTGGAACTTTCCAAGAGTTGAAACTTTATTAATTGAATCAACATACGGTGCA

CENSYa_1545    K  E  D  I  Q  P  T  R  Q  E  V  E  S  A  F  I  N  A  V  N  G  A  L  A  D  G  G  K  V  L
Original 1171 AAAGAGGACATACAGCCCACGCGCCAGGAGGTAGAATCCGCGTTCATCAACGCGGTAAACGGGGCTCTTGCTGACGGCGGCAAGGTCCTC
Optimized     AAAGAAGATATCCAACCAACTAGACAAGAAGTTGAATCAGCGTTTATTAATGCTGTTAATGGAGCATTAGCAGATGGAGGTAAAGTTTTA

CENSYa_1545    I  P  I  P  A  V  G  R  A  Q  E  I  M  M  V  I  D  H  Y  M  K  S  G  E  M  A  E  A  P  V
Original 1261 ATACCGATACCGGCGGTGGGCCGCGCCCAGGAGATAATGATGGTGATCGACCACTATATGAAGTCCGGCGAGATGGCAGAAGCGCCGGTA
Optimized     ATACCTATTCCAGCAGTTGGAAGAGCACAAGAAATAATGATGGTAATTGATCATTACATGAAATCTGGTGAAATGGCTGAAGCACCTGTT

CENSYa_1545    F  T  E  G  M  I  S  E  A  S  A  I  H  E  A  H  P  E  Y  L  A  R  E  L  K  Q  K  I  L  E
Original 1351 TTCACAGAAGGGATGATCTCCGAGGCATCAGCGATACACGAGGCGCACCCCGAGTATCTTGCGCGGGAGCTCAAGCAGAAAATACTCGAG
Optimized     TTTACTGAAGGAATGATTTCTGAAGCTAGTGCTATTCATGAAGCACACCCTGAATATCTTGCTAGAGAACTTAAACAAAAAATTCTTGAA

CENSYa_1545    T  D  D  N  P  F  D  S  E  Y  F  T  N  V  E  H  A  D  G  R  D  E  A  L  R  D  G  S  P  C
Original 1441 ACCGACGACAACCCGTTTGATTCAGAGTACTTTACCAATGTCGAGCATGCTGACGGCAGGGACGAGGCCCTCCGCGACGGCTCGCCGTGC
Optimized     ACTGATGATAATCCTTTTGATTCTGAATATTTTACAAATGTAGAACACGCTGATGGTAGGGATGAAGCATTAAGAGATGGATCACCATGT

CENSYa_1545    I  I  L  A  T  S  G  M  L  E  G  G  P  V  L  E  Y  F  K  S  I  A  P  H  K  Q  N  K  I  L
Original 1531 ATAATACTGGCCACTTCGGGCATGCTCGAGGGCGGGCCCGTCCTGGAGTACTTTAAGAGCATAGCGCCGCACAAGCAGAACAAGATACTT
Optimized     ATTATACTAGCAACTAGTGGAATGTTAGAAGGAGGACCAGTATTAGAATATTTTAAAAGTATTGCACCTCATAAACAAAACAAAATTTTA

CENSYa_1545    F  V  S  Y  Q  V  N  G  T  L  G  R  R  V  L  D  G  A  R  Q  V  P  L  M  N  R  G  G  K  I
Original 1621 TTTGTGTCATACCAGGTCAATGGCACGCTCGGCAGGAGGGTTCTAGACGGGGCGCGGCAGGTCCCGCTTATGAACAGGGGCGGCAAGATA
Optimized     TTTGTTAGTTATCAAGTTAATGGAACTTTAGGTAGAAGAGTTTTAGATGGAGCAAGACAAGTACCTTTAATGAATAGAGGAGGTAAAATT

CENSYa_1545    E  V  V  N  I  E  C  R  M  E  K  L  D  G  F  S  G  H  S  D  Y  N  Q  L  T  G  F  V  Q  K
Original 1711 GAGGTGGTCAACATAGAGTGCAGGATGGAGAAGCTAGACGGGTTCAGCGGCCACAGCGACTACAACCAGCTGACCGGGTTTGTGCAAAAG
Optimized     GAAGTAGTTAATATTGAATGTAGAATGGAAAAATTAGATGGATTTTCAGGACACTCTGATTATAATCAGTTAACTGGATTTGTTCAAAAA

CENSYa_1545    L  R  P  K  L  R  R  V  L  V  N  H  G  E  R  R  K  S  E  N  L  A  L  A  V  R  R  M  F  R
Original 1801 CTCCGGCCCAAGCTGAGGCGAGTCCTTGTAAACCACGGCGAGCGCCGCAAGTCGGAGAACCTTGCGCTTGCGGTGCGCAGGATGTTCAGA
Optimized     TTAAGACCAAAATTAAGAAGAGTATTAGTTAATCACGGAGAAAGAAGAAAAAGTGAAAATCTTGCATTAGCAGTTAGAAGAATGTTTAGG

CENSYa_1545    I  P  A  H  Y  P  Q  I  Q  E  S  I  K  L  F  *
Original 1891 ATACCGGCGCACTATCCGCAGATACAAGAGAGCATAAAGCTGTTCTAG
Optimized     ATTCCAGCACACTATCCACAGATTCAGGAATCTATAAAATTATTTTAA
```

**Supplementary Figure 9. Codon optimized sequences of the *aCPSF1* genes from *Lokiarchaeota* (*Loki-aCPSF1*: *Lokiarch_44440)* and *Thaumarcheae* (*Csy-aCPSF1*: *CENSYa_1545*).** Original, the original protein coding sequences; Optimized, the protein encoding sequences are optimized to compatible to the codon usage basis of *M. maripaludis*.

### Supplementary Tables

#### Table 1. Strains and primers used in this study

| Strains and plasmids | Characteristics and descriptions | Reference or sources |
|---|---|---|
| **Strains** | | |
| *E. coli* DH5α | F$^-$φ80d *lac*ZΔM15 Δ(*lac*ZYA-*arg* F) U169 *end*A1 *rec*A1 *hsd*R17(r$_k^-$,m$_k^+$) *sup*E44λ- *thi* -1 *gyr*A96 *rel*A1 *pho*A | TransGen, Beijing |
| *E. coli* BL21(DE3)pLysS | F$^-$ *omp*T *hsd*S(r$_B^-$m$_B^-$) *gal dcm*(DE3)pLysS Cam$^r$ | TransGen, Beijing |
| *E. coli* BW25113 | *lacI*$^q$ *rrnB3* Δ*lacZ4787 hsdR514* DE(*araBAD*)*567* DE(*rhaBAD*)*568 rph-1* | provided by Pro.Tao |
| *M. maripaludis* S2 (*Mmp*) | Wild-type *M. maripaludis*, Pur$^S$, Neo$^S$ | [1] |
| *Mmp-tetO-aCPSF1* | *hpt*::P*mcr-tetR*-T*mcr*-P*nif-tetO*-His$_6$-*MMP0694*, S2 with *MMP0694* inducible expression | This study |
| *Mmp*-∇*aCPSF1* | *hpt*::P*mcr-tetR*-T*mcr*-P*nif-tetO*-His$_6$-*MMP0694*, *MMP0694*::*pac*, Pur$^R$, strain *tetO-aCPSF1* with the indigenous *MMP0694* deletion | This study |
| *Mmp*-com(*Mmp*-C1) | *hpt*::P*mcr-tetR*-T*mcr*-P*nif-tetO*-His$_6$-*MMP0694*, *MMP0694*::*pac*, pMEV2-*MMP0694*, Pur$^R$, Neo$^R$, ∇*aCPSF1* with *MMP0694* complement | This study |
| *Mmp*-com(*Mmp*-C1mu) | *hpt*::P*mcr-tetR*-T*mcr*-P*nif-tetO*-His$_6$-*MMP0694*, *MMP0694*::*pac*, pMEV2-*MMP0694*H243/246A, Pur$^R$, Neo$^R$, ∇*aCPSF1* with *MMP0694*H243/246A complement | This study |
| *Mmp*-com(*Loki*-C1) | *hpt*::P*mcr-tetR*-T*mcr*-P*nif-tetO*-His$_6$-*MMP0694*, *MMP0694*::*pac*, pMEV2-*Lokiarch44440*, Pur$^R$, Neo$^R$, ∇*aCPSF1* with *Lokiarch44440* complement | This study |
| *Mmp*-com(*Csy*-C1) | *hpt*::P*mcr-tetR*-T*mcr*-P*nif-tetO*-His$_6$-*MMP0694*, *MMP0694*::*pac*, pMEV2-*CENSYa1545*, Pur$^R$, Neo$^R$, ∇*aCPSF1* with *CENSYa1545* complement | This study |
| *Mmp*-S2-pMEV2 | pMEV2, Neo$^R$, S2 with pMEV2 | This study |
| *Mmp*-aCPSF1-F | Φ(*MMP0694*-3Flag), Neo$^R$, S2 with 3Flag tags fused in the C-terminal of *MMP0694* | This study |
| *Mmp*-aRpoL-HF | Φ(MMP0261-His$_6$-3Flag ), Neo$^R$, S2 with His$_6$-3Flag tags fused in the C-terminal of *MMP0261* | This study |
| *Mmp*-aCPSF1-mMaple | Φ(*MMP0694-mMaple3*), Pur$^R$, S2 with *mMaple3* fused in the C-terminal of *MMP0694* | This study |
| *Mmp*-aRpoL-mMaple | Φ(*MMP0261-mMaple3*), Neo$^R$, S2 with *mMaple3* fused in the C-terminal of *MMP0261* | This study |
| *Mmp-*Δ*earA* | *MMP1718*::*neo*, Neo$^R$, S2 with *MMP1718* deletion | This study |
| *Mmp-*∇*aCPSF1* Δ*earA* | *hpt*::P*mcr-tetR*-T*mcr*-P*nif-tetO*-His$_6$-*MMP0694*, *MMP0694*::*pac*, *MMP1718*::*neo*, Pur$^R$, Neo$^R$, ∇*aCPSF1* with *MMP1718* deletion | This study |
| **Plasmids** | | |
| pMD19-T | Amp$^R$ | Takara, Japan |
| pMD19-T-*hpt*up | pMD19-T with *M. maripaludis* S2 *MMP0145* upstream fragment (*hpt*up) inserted into T-overhang end, Amp$^R$ | This study |
| pMD19-T-P*nif-tetO*-His$_6$-*aCPSF1-hpt*down | pMD19-T with P*nif-tetO*-His$_6$-*MMP0694-hpt*down fragment inserted into T-overhang end, Amp$^R$ | This study |
| pMD19-T-P*mcr-tetR*-T*mcr* | pMD19-T with P*mcr-tetR*-T*mcr* fragment inserted into T-overhang ends, Amp$^R$ | This study |
| p*tetR*/*tetO*-His$_6$-*aCPSF1* | pMD19-T-*hpt*up with P*mcr-tetR*-T*mcr* and P*nif-tetO*-His$_6$-*MMP0694-hpt*down fragment inserted into 3′-ends of *hpt*up, Amp$^R$ | This study |
| pMD19-T-*aCPSF1*up | pMD19-T with *M. maripaludis* S2 *MMP0694* upstream fragment (*aCPSF1*up) inserted into T-overhang ends, Amp$^R$ | This study |
| pMD19-T-Δ*aCPSF1* | pMD19-T-*aCPSF1*up with the *pac* gene and *MMP0694* downstream fragment (*aCPSF1*down) inserted into 3′-ends of *aCPSF1*up, Amp$^R$ | This study |
| pIJA03 | Amp$^R$, Pur$^R$ | [2] |
| pMEV2 | Amp$^R$, Neo$^R$ | [2] |
| pMEV2-*MMP0694* | pMEV2 with *MMP0695* promoter, 5′ UTR and ORF of | This study |

| | | |
|---|---|---|
| | *MMP0694* inserted between *Xho* I and *Bgl* II, Amp$^{R}$, Neo$^{R}$ | |
| pMEV2-*MMP0694*H243/246A | pMEV2 with *MMP0695* promoter, 5′ UTR of *MMP0694* and *MMP0694*H243/246A inserted between *Xho* I and *Bgl* II, Amp$^{R}$, Neo$^{R}$ | This study |
| pMEV2-*Lokiarch44440* | pMEV2 with *MMP0695* promoter, 5′ UTR of *MMP0694* and *Lokiarch44440* inserted between *Xho* I and *Bgl* II, Amp$^{R}$, Neo$^{R}$ | This study |
| pMEV2-*CENSYa1545* | pMEV2 with *MMP0695* promoter, 5′ UTR of *MMP0694* and *CENSYa1545* inserted between *Xho* I and *Bgl* II, Amp$^{R}$, Neo$^{R}$ | This study |
| pMD19-T-*aCPSF1*-3Flag | pMD19-T with *MMP0694* 3′-end fragment and 3Flag tag inserted into T-overhang ends, Amp$^{R}$ | This study |
| pMD19-T-*aCPSF1*-3Flag-*pac* | pMD19-T-*aCPSF1*-3Flag with P*mcr*-*pac*-T*mcr* and *MMP0694* downstream fragment inserted into 3′-ends of *aCPSF1*-3Flag, Amp$^{R}$ | This study |
| pMD19-T-*rpoL*-3Flag-His$_6$ | pMD19-T with *MMP0261* 3′-end fragment, 3Flag and His$_6$ tags inserted into T-overhang ends, Amp$^{R}$ | This study |
| pMD19-T-*rpoL*-3Flag-His$_6$-*neo* | pMD19-T-*rpoL*-3Flag-His$_6$ with P*mcr*-*neo*-T*mcr* and *MMP0261* downstream fragment inserted into 3′-ends of *rpoL*-3Flag-His$_6$, Amp$^{R}$ | This study |
| pMD19T-*aCPSF1*-*mMaple3*-*pac* | pMD19-T-*aCPSF1*-3Flag-*pac* with *mMaple3* gene replaced the 3Flag tag, Amp$^{R}$ | This study |
| pMD19-T-*rpoL*-*mMaple3*-*neo* | pMD19-T-*rpoL*-3Flag-His$_6$-*neo* with *mMaple3* gene replaced the 3Flag-His$_6$ tags, Amp$^{R}$ | This study |
| pMD19-T-*earA*up | pMD19-T with *M. maripaludis* S2 *MMP1718* upstream (*earA*up) fragment inserted into T-overhang ends, Amp$^{R}$ | This study |
| pMD19-T-Δ*earA* | pMD19-T-*earA*up with the *neo* gene and *MMP1718* downstream (*earA*down) fragment inserted into 3′-ends of *earA*up, Amp$^{R}$ | This study |
| pSB1s | Str$^{R}$ | provided by Prof.Tao |
| pSB1s-His$_6$-SUMO-*aCPSF1* | pSB1s with His$_6$-SUMO and *MMP0694* genes inserted into the expression region, Str$^{R}$ | This study |
| pGEX-4T-1 | Amp$^{R}$ | |
| pGEX-4T-1-*rpoD* | pGEX-4T-1 with *M. maripaludis* S2 *MMP1322* inserted into *Eco*R I, Amp$^{R}$ | This study |
| pGEX-4T-1-*rpoD/L* | pGEX-4T-1-*rpoD* with the sequence from RBS to stop codon of *M. maripaludis* S2 *MMP0261* inserted into *Xho* I, Amp$^{R}$ | This study |

**Table 2. Primers used in this study.**

| Primer | Sequence (5′-3′) | Purpose |
|---|---|---|
| *Hpt*up-F | TGTCGGGGGAGTTCAGTCC | Construction of strain ∇*aCPSF1* |
| *Hpt*up-R | ATGTTTTCAATGATTCTTCCAATAATT | Construction of strain ∇*aCPSF1* |
| P*mcr*-F | GGAAGAATCATTGAAAACATGGATGATTAATTTTAAGAGA | Construction of strain ∇*aCPSF1* |
| P*mcr*-R | TTTATCTAATCTAGACATCATGAGAATCACTCCTATTTTT | Construction of strain ∇*aCPSF1* |
| *TetR*-F | AAAAAATAGGAGTGATTCTCATGATGTCTAGATTAGATAAA | Construction of strain ∇*aCPSF1* |
| *TetR*-R | GGGTCGTGGGGCGGGCGTTAAGACCCACTTTCACA | Construction of strain ∇*aCPSF1* |
| T*mcr*-F | TGTGAAAGTGGGTCTTAACGCCCGCCCCACGACCC | Construction of strain ∇*aCPSF1* |
| T*mcr*-R | CTATATAAAGTTTTCGCCCCTATCAACCCAGTGAATTAAAATATAT | Construction of strain ∇*aCPSF1* |
| P*nif-tetO*-His$_6$-*aCPSF1*-F | TTGATAGGGGCGAAAACTTTATATAGCCCTATCAGTGATAGAGAGTTCACAACAATATATAGAGGCCTAAAAAATGCATCATCATCATCATCACTCAGCTGAAGATATATTAAACGA | Construction of strain ∇*aCPSF1* |
| P*nif-tetO*-His$_6$-*aCPSF1*-R | GTTTACTTTTCCGTCAACAACTTCATTATCTCAATCTTATTGAATC | Construction of strain ∇*aCPSF1* |
| *Hpt*down-F | ATAAGATTGAGATAATGAAGTTGTTGACGGAAAAGTAA | Construction of strain ∇*aCPSF1* |
| *Hpt*down-R | TCCAATGGTTCCCCCTAATC | Construction of strain ∇*aCPSF1* |
| 19T*hpt/aC1*up-F | GCATGCAAGCTTGGCGTA | Construction of strain ∇*aCPSF1* |
| 19T*hpt*up-R | AATTATTGGAAGAATCATTGAAAACAT | Construction of strain ∇*aCPSF1* |
| *aCPSF1*up-F | GATTAATATTAAGTGGTGATAAGATGAT | Construction of strain ∇*aCPSF1* |
| *aCPSF1*up-R | AATAGTCCCTCCTAATAATATGTCTG | Construction of strain ∇*aCPSF1* |
| 19T*aC1*up-R | AATAGTCCCTCCTAATAATATGTCTGTTTAAATTC | Construction of strain ∇*aCPSF1* |
| *Pac*-(*aC1*up)-F | TTATTAGGAGGGACTATTATGACCGAGTACAAGCCCACG | Construction of strain ∇*aCPSF1* |
| *Pac*-(*aC1*dw)-R | AAAAGAATACTCAGGCACCGGGCTTGCG | Construction of strain ∇*aCPSF1* |
| *aCPSF1*dw-F | GTGCCTGAGTATTCTTTTTTATTTTTAATTCGAATAATAGATATAC | Construction of strain ∇*aCPSF1* |
| *aCPSF1*dw-R | GCCAAGCTTGCATGCGTTCTGATTTCTTATGGAATTATTTATC | Construction of strain ∇*aCPSF1* |
| *aCPSF1Xho*I-F | CCGCTCGAGCCGGTGAGTATATAAAGCATTTATTATAAATTGATTAATATTAGGAGGGACTATTATGTCAGCTGAAGATATATTAAACG | Construction of complementary strain |
| *aCPSF1Bgl*II-R | GGAAGATCTTTATCTCAATCTTATTGAATCGAGA | Construction of complementary strain |
| pMEV2-(*Loki*C1)-F | AGATCTCATGATATCTAGATCC | Construction of complementary strain |
| pMEV2-(*Loki*C1)-R | CTCGAGCTCCCTGAAGAAG | Construction of complementary strain |
| *Loki*C1-(pMEV2)-F | TCTCTTCTTCTTCAGGGAGCTCGAGCCGGTGAGTATATAAAG | Construction of complementary strain |
| *Loki*C1-(pMEV2)-R | CTAGAGGATCTAGATATCATGAGATCTTTATTTTAAAAGAACTACTTCTAAGTTTTTAG | Construction of complementary strain |
| *Cys*C1-(pMEV2)-F | TCTCTTCTTCTTCAGGGAGCTCGAGCCGGTGAGTATATAAAG | Construction of complementary strain |
| *Csy*C1-(pMEV2)-R | TCCTCTAGAGGATCTAGATATCATGAGATCTTTAAAATAATTTTATAGATTCCTGAATC | Construction of complementary strain |
| *aCPSF1*-H243A-F | GTAGTAACTCACGCGGCCCTTGACCACTGTG | Site-directed mutation of *aCPSF1* |
| *aCPSF1*-H243A-R | CACAGTGGTCAAGGGCCGCGTGAGTTACTAC | Site-directed mutation of *aCPSF1* |
| *aCPSF1*-H246A-F | CGCGGCCCTTGACGCCTGTGGATTTATTC | Site-directed mutation of *aCPSF1* |
| *aCPSF1*-H246A-R | GAATAAATCCACAGGCGTCAAGGGCCGCG | Site-directed mutation of *aCPSF1* |
| *EarA*dw-F | AATCATGGCTTTTAATATAGATGGTTCG | Deletion of *earA* |
| *EarA*dw-R | TTAGTGTGTGGATGCATAACACG | Deletion of *earA* |
| 19T*earA*dw-F | AATCATGGCTTTTAATATAGATGG | Deletion of *earA* |
| 19T*earA*dw-R | TCTAGAGGATCCCCGGGTAC | Deletion of *earA* |
| *EarA*up-F | GTACCCGGGGATCCTCTAGAATGTAGATATATTTTATAC | Deletion of *earA* |

| | | |
|---|---|---|
| | GGAAGGTTC | |
| *EarA*up-R | GTTCAATCATAAAAGACACCTCGAAAGTTAAAAATTAATTAAAATTAC | Deletion of *earA* |
| *Neo*-(*earA*up)-F | GGTGTCTTTTATGATTGAACAAGATGGATTG | Deletion of *earA* |
| *Neo*-(*earA*dw)-R | CTATATTAAAAGCCATGATTTCAGAAGAACTCGTCAAG | Deletion of *earA* |
| *aCPSF1*Flag-F | CACAGGAGACATTAAATTTGAAG | Construction of strain aCFSF1-F |
| *aCPSF1*Flag-R | TTATGCATAATCTGGAACATCATATGGATAAACTTTTAATTTGTCATCGTCATCTTTATAATCTTTGTCATCGTCATCTTTATAATCTTTGTCATCGTCATCTTTATAATCGATAGATCTCAATCTTATTGAATCGAG | Construction of strain aCFSF1-F |
| *Pac*-(*C1*F)-F | CCAGATTATGCATAAAATCGAAAGGAAACCTAATATGGTTTC | Construction of strain aCFSF1-F |
| *Pac*-(*C1*dw)-R | CCGATATATCTTATGCTCCTGGTTTTCTTG | Construction of strain aCFSF1-F |
| *aCPSF1*Flagdw-F | AGGAGCATAAGATATATCGGGGTGATACAATG | Construction of strain aCFSF1-F |
| *aCPSF1*Flagdw-R | CAAGCTTGCATGCCTGCAGGCGTTTTAAAGGCAATAAAATAGTATTATTG | Construction of strain aCFSF1-F |
| *RpoL*Flag-F | CATTGATTATTCATTTGACGCA | Construction of strain aRpoL-HF |
| *RpoL*Flag-R | TTAGTGGTGGTGGTGGTGGTGCTTATCGTCATCGTCCTTGTAGTCCTTATCGTCATCGTCCTTGTAGTCCTTATCGTCATCGTCCTTGTAGTCTAAATCTTCAAGGGTTTTGTTACAAAG | Construction of strain aRpoL-HF |
| 19T*rpoL*Flag-F | CATGGTCATAGCTGTTTC | Construction of strain aRpoL-HF |
| 19T*rpoL*Flag-R | CTGCAGGTCGACGATTTTAG | Construction of strain aRpoL-HF |
| *Neo*-(*rpoL*F)-F | CTAAAATCGTCGACCTGCAGATAAAAAACGCCCTATTCG | Construction of strain aRpoL-HF |
| *Neo*-(*rpoL*dw)-R | TATAAAATATCAATCAGAAGAACTCGTCAAG | Construction of strain aRpoL-HF |
| *RpoL*Flagdw-F | GTTCTTCTGATTGATATTTTATATCGGCATAATTTATTTTTTTAAC | Construction of strain aRpoL-HF |
| *RpoL*Flagdw-R | AGGAAACAGCTATGACCATGCCCATTGTAAGTTTTGTAAATG | Construction of strain aRpoL-HF |
| 19T-*aCFSF1*F-F | GAAAGGAAACCTAATATGGTTTCC | Construction of strain aCPSF1-mMaple |
| 19T-*aCFSF1*F-R | ATCGATAGATCTCAATCTTATTGAATC | Construction of strain aCPSF1-mMaple |
| mMaple-(*aC1*)-F | TAAGATTGAGATCTATCGATATGGTTTCAAAAGGAGAAG | Construction of strain aCPSF1-mMaple |
| mMaple-(*aC1*)-R | ACCATATTAGGTTTCCTTTCTTATTTGTATAATTCATCCATACCAC | Construction of strain aCPSF1-mMaple |
| 19T-*rpoL*F-F | CGTCGACATAAAAAACGC | Construction of strain aRpoL-mMaple |
| 19T-*rpoL*F-R | TAAATCTTCAAGGGTTTTGTTACAAAGATC | Construction of strain aRpoL-mMaple |
| mMaple-(*rpoL*)-F | ACAAAACCCTTGAAGATTTAATGGTTTCAAAAGGAGAAG | Construction of strain aRpoL-mMaple |
| mMaple-(*rpoL*)-R | GGGCGTTTTTTATGTCGACGTTATTTGTATAATTCATCCATACCAC | Construction of strain aRpoL-mMaple |
| pSB1s-F | TACAGATTAAATCAGAACGCAG | Protein expression of *Mmp*-aCPSF1 |
| pSB1s-R | GGTTAATTCCTCCTGTTAGC | Protein expression of *Mmp*-aCPSF1 |
| SUMO-(pSB1s)-F | GCTAACAGGAGGAATTAACCATGGGCAGCAGCCATCAT | Protein expression of *Mmp*-aCPSF1 |
| SUMO-(*aCPSF1*)-R | CTTCAGCTGACATACCACCAATCTGTTCTCTG | Protein expression of *Mmp*-aCPSF1 |
| *aCPSF1*-(SUMO)-F | CAGATTGGTGGTATGTCAGCTGAAGATATATTAAACG | Protein expression of *Mmp*-aCPSF1 |
| *aCPSF1*-(pSB1s)-R | GCGTTCTGATTTAATCTGTATTATCTCAATCTTATTGAATCGAGATTCATTG | Protein expression of *Mmp*-aCPSF1 |
| *RpoD*-*Eco*RI-F | GGAATTCATGAAAATGGAATTAAAAGCCCC | Protein expression of *Mmp*-aRpoD/L |
| *RpoD*-*Eco*RI-R | GGAATTCTTAATTTTCGTCACTTAATTCTTGGC | Protein expression of *Mmp*-aRpoD/L |
| *RpoL*-*Xho*I-F | CCGCTCGAGGAGGTAATTAAATATGAACTACGTTAAAATCATTG | Protein expression of *Mmp*-aRpoD/L |
| *RpoL*-*Xho*I-R | CCGCTCGAGTTATAAATCTTCAAGGGTTTTGTTACA | Protein expression of *Mmp*-aRpoD/L |
| P1719-NB-F | TGGGGAAACCTACGATATAGTC | Northern Blot |
| P1719-NB-R | GCCCCTATTTTCTTCTTCAGTAGC | Northern Blot |
| P1718-NB-F | TTGGGATCAGAAGACATGATATTT | Northern Blot |
| P1718-NB-R | ACCAGTTATCGGCAGAATCGTA | Northern Blot |
| P1717-NB-F | GCAGATTCTTTGAAATTATTCCTCA | Northern Blot |
| P1717-NB-R | TTCGACTACGATACAGTTATCTCCA | Northern Blot |
| PflaB2-NB-F | TTGGTACCTTGATTGTTTTTATTGC | Northern Blot |

| | | |
|---|---|---|
| PflaB2-NB-R | TAGTGTTGTAAAGGTCTCCACCTG | Northern Blot |
| P1099-NB-F | TCATATCCCTGCAAAGCACT | Northern Blot |
| P1099-NB-R | GGAGATATGGCTTCAAGAATTGC | Northern Blot |
| P1100-NB-F | CAACAGTTCGTGAATCGATACTCG | Northern Blot |
| P1100-NB-R | TTTTACAGGAACGAACGGGA | Northern Blot |
| P1146-NB-F | CCAGTTCAATTTAGTTATTCTATCGA | Northern Blot |
| P1146-NB-R | TCCTATTTCTTTACCGTTCCC | Northern Blot |
| P1147-NB-F | CACGGTGAAGTAAATGACAAAAG | Northern Blot |
| P1147-NB-R | CTCTGTTTCCTTTTCCGGTC | Northern Blot |
| P1149-NB-F | CATGCTTGAAGCTATTGATGAAGG | Northern Blot |
| P1149-NB-R | CGATGCAAGGTACACGTC | Northern Blot |
| P1150-NB-F | CTTCATTTACCCCATAAGATCTTGC | Northern Blot |
| P1150-NB-R | CTTGGAAACGTAAGTCCTTCAGA | Northern Blot |
| P0155-NB-F | GTGCACTTGAAAATGGCGA | Northern Blot |
| P0155-NB-R | CTCTTAAGGAGTGCTTTTTAGACC | Northern Blot |
| 3′R-RT-P | ATTGATGGTGCCTACAG | 3′RACE |
| 1100-3′R-N1-F | TGGATAAAGAGGGCTTGGG | 3′RACE |
| 1100-3′R-N2-F | GAATGATTTGATAATGGTTGATAGACGG | 3′RACE |
| 1147-3′R-N1-F | GGAAGAAGAGCATTCCACGTAA | 3′RACE |
| 1147-3′R-N2-F | TATGTGCAGCATGCGGATTT | 3′RACE |
| 1149-3′R-N1-F | CATGCTTGAAGCTATTGATGAAGG | 3′RACE |
| 1149-3′R-N2-F | CCATCATGTTCTGCATGTATGG | 3′RACE |
| 0155-3′R-N1-F | AAAGCAGAAATGGCCGAAA | 3′RACE |
| 0155-3′R-N2-F | CAGCGTAGTTGGAAATGTTATCAC | 3′RACE |
| 0900-3′R-N1-F | GATGGAATAAAACGAATAGTTTCAG | 3′RACE |
| 0900-3′R-N2-F | GAAGCCAATCAATTGCAACG | 3′RACE |
| 0901-3′R-N1-F | CGTTCCAATTTTAGTTGCACTTAC | 3′RACE |
| 0901-3′R-N2-F | GAAGAAGATCTTGGAGAATATGAAGT | 3′RACE |
| 1241-3′R-N1-F | CACAAACAGCTACCCCGAGA | 3′RACE |
| 1241-3′R-N2-F | TGCCCAAACTGGTGAAAAATAC | 3′RACE |
| 1242-3′R-N1-F | GAACTGACGCAATCTGCAAA | 3′RACE |
| 1242-3′R-N2-F | CGATGAAAAACATAACTTTAGAAAAAATAG | 3′RACE |
| 1640-3′R-N1-F | CATCTTGTCAAACATTATTGCAGAAGA | 3′RACE |
| 1640-3′R-N2-F | GAAGGAGTAAGAGAATGTCAGATCA | 3′RACE |
| 1641-3′R-N1-F | CAGGAAGGATATACTGTAGTAATG | 3′RACE |
| 1641-3′R-N2-F | GGGGATGCTTCAAACGATATT | 3′RACE |
| 0291-3′R-N1-F | GGTGCTAAACAGATTGGGGT | 3′RACE |
| 0291-3′R-N2-F | CAGGGAAGCTGCATACACA | 3′RACE |
| 0292-3′R-N1-F | CGTTGGGATTCAGCTGTCT | 3′RACE |
| 0292-3′R-N2-F | GATACTCGGACTTATTGATACTCTTGA | 3′RACE |
| 1648-3′R-N1-F | GATATTGAATCACTATTTGGTGGA | 3′RACE |
| 1648-3′R-N2-F | GTTACGGTAATGGTTGGCC | 3′RACE |
| 1661-3′R-N1-F | GGTGAAAATGATGGAAAATAGAGA | 3′RACE |
| 1661-3′R-N2-F | GCCTTATGTTCAGAATATGGAAT | 3′RACE |
| 1719-3′R-N1-F | AAACTTACGTAGAACCCAAAACCG | 3′RACE |
| 1719-3′R-N2-F | CTACTGAAGAAGAAAATAGGGGCA | 3′RACE |
| 0295-3′R-N1-F | CCAATTTAATTGACGGATATACGGAA | 3′RACE |
| 0295-3′R-N2-F | GACATGGTCTATGGTATAACGATTAGT | 3′RACE |
| 0296-3′R-N1-F | GCACTTGTCTGGAACGAAATC | 3′RACE |
| 0296-3′R-N2-F | CGCCTGATACGATAACTGC | 3′RACE |
| 1016-3′R-N1-F | CTCTACGTTGCAGAAGGAAAAC | 3′RACE |
| 1016-3′R-N2-F | GGCGAAGAAATGGAATTAGTTGC | 3′RACE |
| 1017-3′R-N1-F | GGAGGTATGTGTAGTTTCAGTTGT | 3′RACE |
| 1017-3′R-N2-F | GAGGCAGTTTCAGAAAGCG | 3′RACE |
| 0137-3′R-N1-F | ACATTACAACAGCAGTTCCC | 3′RACE |
| 0137-3′R-N2-F | TGGGATGGGTCATTAAGCG | 3′RACE |
| Term-3′adapter | NN-8mer_index-NNNNAGATCGGAAGAGCGTCGTGT (5′ phosphorylated, 3′ amino blocked) | Term-seq |
| Term-RT-P | TCTACACTCTTTCCCTACACGACGCTCTTC | Term-seq |
| cDNA 3′adapter | GCAGATCGGAAGAGCACACGTCTGAACTCCAGTCAC (5′ phosphorylated, 3′ amino blocked) | Term-seq |
| PCR-forward pri mer | AATGATACGGCGACCACCGAGATCTACACTCTTTCCCT ACACGACGCTCT | Term-seq |
| PCR-reverse pri | CAAGCAGAAGACGGCATACGAGAT-8mer_index-GTGAC | Term-seq |

| | | |
|---|---|---|
| mer | TGGAGTTCAGAC | |
| 1100-ChIP-F | CAGCACCTTCAGAAATCG | ChIP |
| 1100-ChIP-R | CATCAACTACAATTTTTACAGGAAC | ChIP |
| 1149-ChIP-F | CAGAAACTTCAGAGGAAGAGA | ChIP |
| 1149-ChIP-R | GGTTTTAATTAGATTAGAGATCTCTTG | ChIP |
| 0901-ChIP-F | GAAGGATTAGATAAACTTGAAGAAGATCT | ChIP |
| 0901-ChIP-R | CTTCTAAAAGGCCCATTAAACC | ChIP |
| 0791-ChIP-F | ATGAGTTTTAACATGTATGTGCC | ChIP |
| 0791-ChIP-R | TCTTTTGAAGTGGTTTCTTTCAA | ChIP |
| 0136-ChIP-F | CATGTAAAAGAAGGCGACGTT | ChIP |
| 0136-ChIP-R | CTTCAAGTGTGGCATTAATCCG | ChIP |
| 0137-ChIP-F | GCGATATACATTACAACAGCAGTT | ChIP |
| 0137-ChIP-R | CTCCATAAACGAGCATTGGAAAT | ChIP |
| 0180-ChIP-F | GCAGTGAACGGAAAAGAATTAGA | ChIP |
| 0180-ChIP-R | CTTGCAGCTCCTGCATC | ChIP |
| 0181-ChIP-F | CATTGACATTGAAAAACGGTGG | ChIP |
| 0181-ChIP-R | GGAAGTTCAAAATATCTGAACTTG | ChIP |
| 0183-ChIP-F | GGACTCACCCCTATTACG | ChIP |
| 0183-ChIP-R | GAGAATTCTTTGAGGTAGTATTCAA | ChIP |
| 0195-ChIP-F | GGAAAGTATTGAAACGTTATCCCTTG | ChIP |
| 0195-ChIP-R | CGTGCGACATTTTTTCATTTGT | ChIP |
| 0210-ChIP-F | CATGGGGCTAGTAAAACAATGT | ChIP |
| 0210-ChIP-R | GGTTGAATTTATGTATACTCCGCC | ChIP |
| 0309-ChIP-F | CGCAATTGAAGCAGGAGC | ChIP |
| 0309-ChIP-R | CTGCTTCTTTTACGAAAGCTATTAAAT | ChIP |
